# The primary sclerosing cholangitis and ulcerative colitis colonic mucosa defined through paired microbial and single-cell RNA sequencing

**DOI:** 10.1101/2024.08.12.607536

**Authors:** Jacqueline LE Tearle, Fan Zhang, Katherine JL Jackson, Pratibha Malhotra, Paris Tavakoli, Sabrina Koentgen, Joanna Warren, Cameron Williams, Ashraful Haque, Arteen Arzivian, Nicodemus Tedla, Andrew Kim, Hamish W King, Georgina L Hold, Simon Ghaly, Kylie R James

## Abstract

Primary sclerosing cholangitis (PSC) is a chronic progressing cholestatic disease that often co-occurs with inflammatory bowel disease (PSC-IBD). PSC-IBD affecting the colon (PSC-UC) is likened clinically to ulcerative colitis (UC), however differences include a right colon dominance, less severe inflammatory presentation and a greater lifetime risk of colorectal cancer. To understand the basis of clinical differences, we combine single-cell mRNA and antigen receptor sequencing, 16S ribosomal DNA analysis and spatial transcriptomics on biopsies from multiple colon regions of both PSC-UC and UC patients in remission or at the time of relapse. We discover disease-specific cell and microbial profiles between these cohorts, highlighting a distinct landscape in the right colon of PSC-UC patients and an epithelial-endothelial cell state that may contribute to intestinal permeability in UC. We show the expansion of an activated mast cell state in both diseases during flare, and demonstrate the requirement of TMEM176B in sustaining this activated state. Together this work demonstrates that PSC-UC and UC are distinct diseases with common cell mechanisms during inflammation, providing cellular and microbial insights to improve treatment of both patient cohorts.

## Introduction

Primary sclerosing cholangitis (PSC) is a rare and chronic cholestatic disease characterised by stricturing of bile ducts, destruction of the biliary tract and fibrosis. Approximately 70% of PSC individuals have concomitant inflammatory bowel disease (PSC-IBD; approximately 2% of IBD cases)^1,2^. 80% of PSC-IBD resembles ulcerative colitis (UC), with colonic involvement (hereafter referred to PSC-UC), while only 10% of PSC-IBD resembles Crohn’s disease^3^. However, studies comparing PSC-UC and UC clinical presentations highlight their distinctions - PSC-UC is more likely to be pancolitis with a right-sided dominance, feature rectal sparing and backwash ileitis, and has a greater lifetime risk of progressing to colitis-associated colorectal cancer compared to UC. This has led to the postulation of PSC-IBD as a distinct disease entity from IBD^4,5^.

Investigations into PSC-IBD and UC disease mechanisms have pointed to disease-specific intestinal cell signatures^6^, microbiota abundances^7^ and risk-associated genetic variants^8^. However, few studies have assessed the contribution of intestinal cells and microbiota in parallel and none have compared the colonic mucosa in the absence of active inflammation, missing potential co-variance between these factors and baseline differences between these diseases. Furthermore, no published studies to date have compared the full range of immune and non-immune cell types of the PSC-UC and UC colonic mucosa with single-cell expression technologies. One recent study performed single-cell RNA sequencing of sorted T and B cells of the PSC-IBD colon^9^, but did not profile myeloid or innate immune cells, or stromal cell populations, which may be implicated in disease pathology and inflammation, to the same granularity^10^. The cellular and microbial basis for clinical differences between PSC-UC and UC remain poorly defined and are instrumental in guiding disease-specific clinical management.

To comprehensively assess intestinal cells and bacteria in PSC-UC and UC we performed integrated single-cell transcriptomics and bacterial amplicon analysis of colonic mucosa biopsies from varied disease-relevant anatomic regions and inflammation states. Integration with published healthy reference data^11^ revealed disease-specific T cell compositions, with anatomical patterning in PSC-UC that mirrors colon regions most commonly affected by inflammation. We discover a rare UC-specific epithelial-endothelial cell state that may be implicated in the vascularisation of the diseased colon. Finally, we discover a novel *TMEM176B+* mast cell state shared by PSC-UC and UC in the context of active inflammation and show suppression of this state upon therapeutic targeting of TMEM176B.

## Results

### Disease- and region-specific microbial signatures of the UC and PSC-UC gut

We previously demonstrated distinct microbial niches across the healthy human colon, with a greater abundance of *Enterococcus* at the right colon and *Bacteroides*, *Coprobacillus* and *Escherichia/Shigella* at the sigmoid colon^12^. Furthermore, others have reported varying microbial compositions between the PSC-IBD and UC fecal^13^, ileal^14^ and right colonic^14^ microbiomes^7^. Therefore, we compared the mucosa-adherent microbial composition of the ascending, transverse and descending colon and rectum between UC and PSC-UC. This was achieved by performing 16S rRNA gene sequencing on mucosal pinch biopsies collected during routine endoscopies from four regions of four patients for each disease cohort (**Figure 1A**). To contextualise our findings, we pulled in published healthy control data from individuals with no reported intestinal pathology^12^. Relevant clinical information, including inflammatory status, for each patient biopsy can be found in **Supplementary Table 1**.

**Figure 1.**
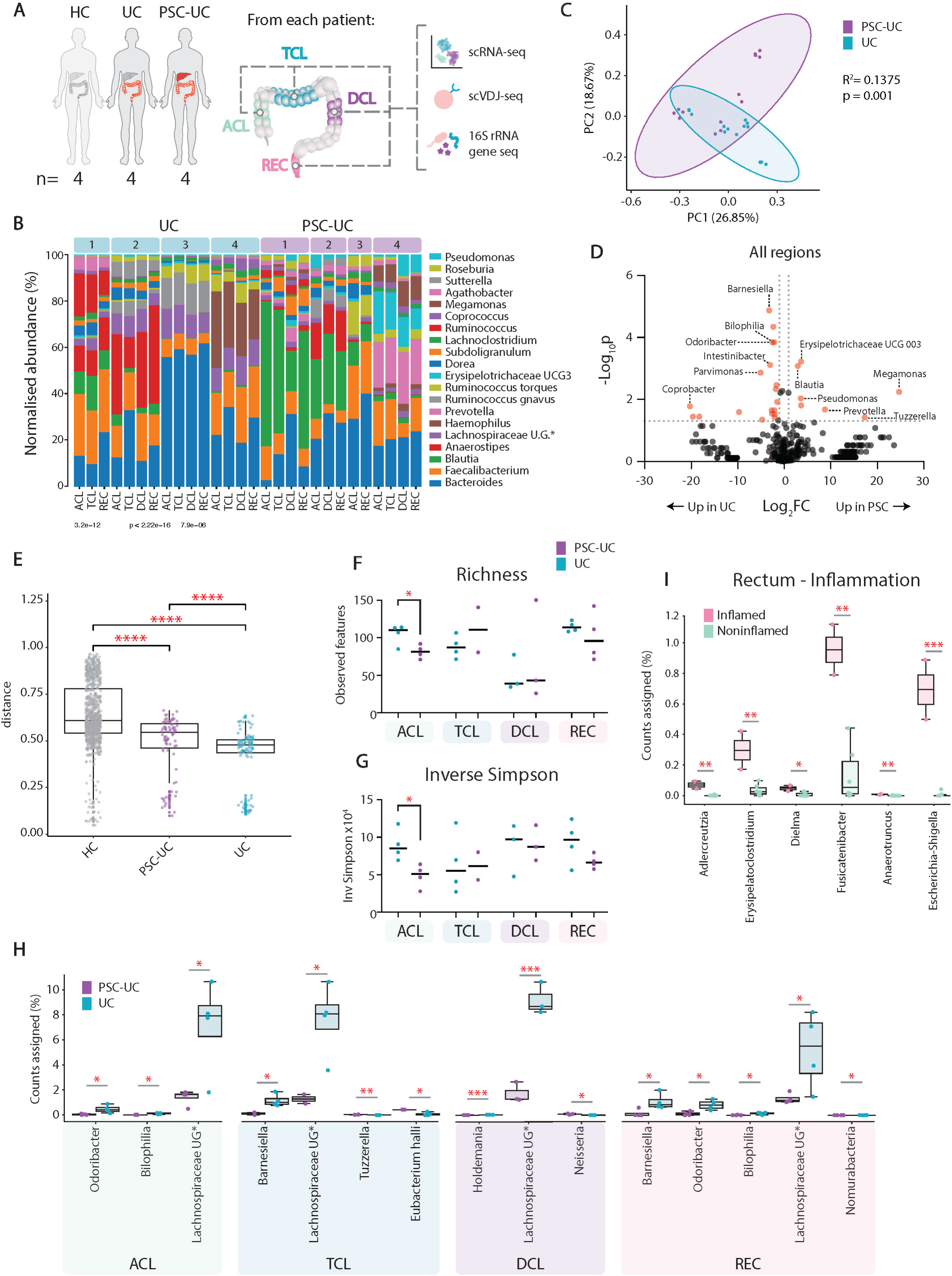
Study overview and microbial analyses. **A** | Study overview. HC, healthy control; UC, ulcerative colitis; PSC-UC, primary sclerosing cholangitis concomitant with colitis (n=4/cohort); ACL, ascending colon; TCL, transverse colon; DCL, descending colon; REC, rectum. Published single-cell RNA sequencing^11^ and 16S rRNA gene sequencing data^12^ of individuals without intestinal disease was used as HC**. B** | Top 20 most abundant genera across all patient samples. Individual patients indicated by blue (UC) and purple (PSC-UC) boxes. **C** | Principle coordinate (PC) analysis plot. Dots represent biopsies. **D** | Volcano plot showing differentially-abundant bacterial genera in UC and PSC-UC patients. Significant genera shown in red and indicated by dotted lines at y = 1.3 (p>0.05) and x = 1, -1. **E** | Bray-Curtis distance between all biopsy combinations for HC, PSC-UC and UC. P values as determined by Wilcoxon Rank Sum for HC vs. PSC-UC = 3.2×10^-12^, for HC vs. UC = 2.22×10^-16^ and for PSC-UC vs UC 7.9×10^-6^. **F** | Richness and **G |** Inverse Simpson index scores of genus counts from 16S rRNA gene sequencing per region. Each dot represents a single biopsy. *P* = 0.0289 for richness and 0.0198 for inverse Simpson for PSC-UC vs. UC in the ACL as determined by unpaired T test. **I** | Percentage of counts assigned to bacterial genus per sample by inflammatory state. Each dot represents one sample (biopsy). P values = 0.0013 (*Adlercreutzia*), 0.0071 (*Erysipelatoclostridium*), 0.0178 (*Dielma*), 0.0019 (*Fusicatenibacter*), 0.0041 (*Anaerotruncus*), 0.0003 (*Escherichia*-*Shigella*), determined by unpaired T test. **H** | Percentage of counts assigned to bacterial genus per sample by region. Each dot represents one sample (biopsy). P values, left to right: 0.0342 (*Odoribacter*), 0.0107 (*Bilophilia*), 0.0241 (*Lachnospiraceae* UG), 0.0458 (*Barnesiella*), 0.0465 (*Lachnospiraceae* UG), 0.0020 (*Tuzzerella*), 0.0205 (*Eubacterium Halli Group*), 0.0005 (*Holdemania*), 0.0010 (*Lachnospiraceae* UG), 0.0346 (*Neisseria*), 0.0370 (*Barnesiella*), 0.0162 (*Odoribacter*), 0.0229 (*Bilophilia*), 0.0469 (*Lachnospiraceae UG*), 0.0403 (*Nomurabacteria*).

Overall, we identified 378 genera across 28 mucosal samples (**Figure 1B**). Principal coordinate analysis revealed that disease type was a major source of variability in bacterial genera (**Figure 1C**) and overall microbial compositions of each anatomical region were largely conserved within each patient, even during inflammation (**Figure 1B**, **Supplementary Figure 1A**). To further compare the PSC-UC and UC mucosal-adherent microbiomes, we compared the relative abundances of each bacterial genera between pooled disease cohorts. This identified a significant enrichment of genera including *Barnesiella*, *Bilophilia*, *Odoribacter*, *Intestinibacter*, *Parvimonas* and *Comprobacter* in UC mucosa compared to PSC-UC mucosa. Fewer genera were significantly enriched in PSC-UC mucosa: Erysipelotrichaceae UCG 003, *Blautia*, *Megamonas*, *Pseudomonas*, *Prevotella*, *Tuzzerella* and *Curvibacter* (**Figure 1D**). PSC-UC and UC patients therefore harbour distinct mucosal adherent intestinal microbiomes.

Following our observation of more enriched genera in UC compared to PSC-UC patients, we asked whether the UC microbiome was more conserved between patients than the PSC-UC microbiome. To this end, we calculated the Bray-Curtis distance between any combination of biopsies for HC, UC and PSC-UC. This showed that UC patients exhibit significantly reduced variation in mucosal microbiome genera compared to both PSC-UC and HC (**Figure 1E**)

We next compared the PSC-UC and UC microbiome in each anatomical region. PSC-UC biopsies displayed reduced bacterial richness (**Figure 1F**) and inverse Simpson (**Figure 1G**) values compared to UC in the ascending colon, the region most commonly affected in PSC-UC. However no difference in Shannon diversity was observed (**Supplementary Figure 1B**). The PSC-UC ascending colon also displayed a reduced abundance of *Odoribacter*, *Bilophilia* and an unclassified genus of the *Lachnospiraceae* family (denoted *Lachnospiraceae* UG) compared to UC (**Figure 1H**, ACL). In the transverse colon, PSC-UC patients displayed a reduction in the abundance of *Barnesiella* and *Lachnospiraceae* UG, and an enrichment of *Tuzzerella* and *Eubacterium hallii* (**Figure 1H**, TCL). Compared to UC, the descending colon of PSC-UC patients exhibited an increased abundance of *Neisseria* and a decreased abundance of *Holdemania* and *Lachnospiraceae UG* compared to UC patients (**Figure 1H**, DCL). Finally, the rectum of PSC-UC patients harboured a significantly lower abundance of *Barnesiella*, *Odoribacter*, *Bilophilia*, *Lachnospiraceae* and *Nomurabacteria* (**Figure 1H**, REC). PSC-UC and UC patients therefore exhibit region-specific dissimilarities in bacterial composition.

We then considered the effect of inflammation on bacterial composition. This analysis was restricted to the rectum, for which our dataset had inflamed biopsies of one PSC-UC and two UC patients. Among enriched genera in inflammation compared to non inflamed tissue were *Adlercreutzia*, *Erysipelatoclostridium*, *Dielma*, *Fusicatenibacter*, *Anaerotruncus* and *Escherichia*-*Shigella* (**Figure 1I**). Overall, we report a unique mucosal microbiome for UC compared to PSC-UC, with the latter having greater similarity to the healthy colon. Therefore, for the PSC-UC disease mechanism in particular, other molecular pathways are likely at play.

### Shared features of antigen presentation in the PSC-UC gut

We examined the cellular landscape of the PSC-UC and UC intestines by performing single-cell gene expression analysis on patient biopsies matched to those used for bacterial analysis (**Methods**, **Figure 1A**). At the time of sampling, one PSC-UC patient and two UC patients displayed active inflammation in one or more regions (**Supplementary Table 1**). Published scRNA-seq data^11^ of colon mucosa resections from individuals with no reported intestinal pathology were integrated to contextualise our findings. Stringent quality control was followed by Leiden clustering and differential gene expression analysis to define broad cell lineages (**Figure 2A-C, Supplementary Figure 1C-D**). Cell lineages were serially subclustered for fine-grain annotation of cell subtypes and states (**Methods**, **Supplementary Figure 1E**). The final dataset consisted of 156,331 high-quality cells across 62 cell types or states, including 51,092 from PSC-UC patients and 27,291 cells from UC patients.

**Figure 2.**
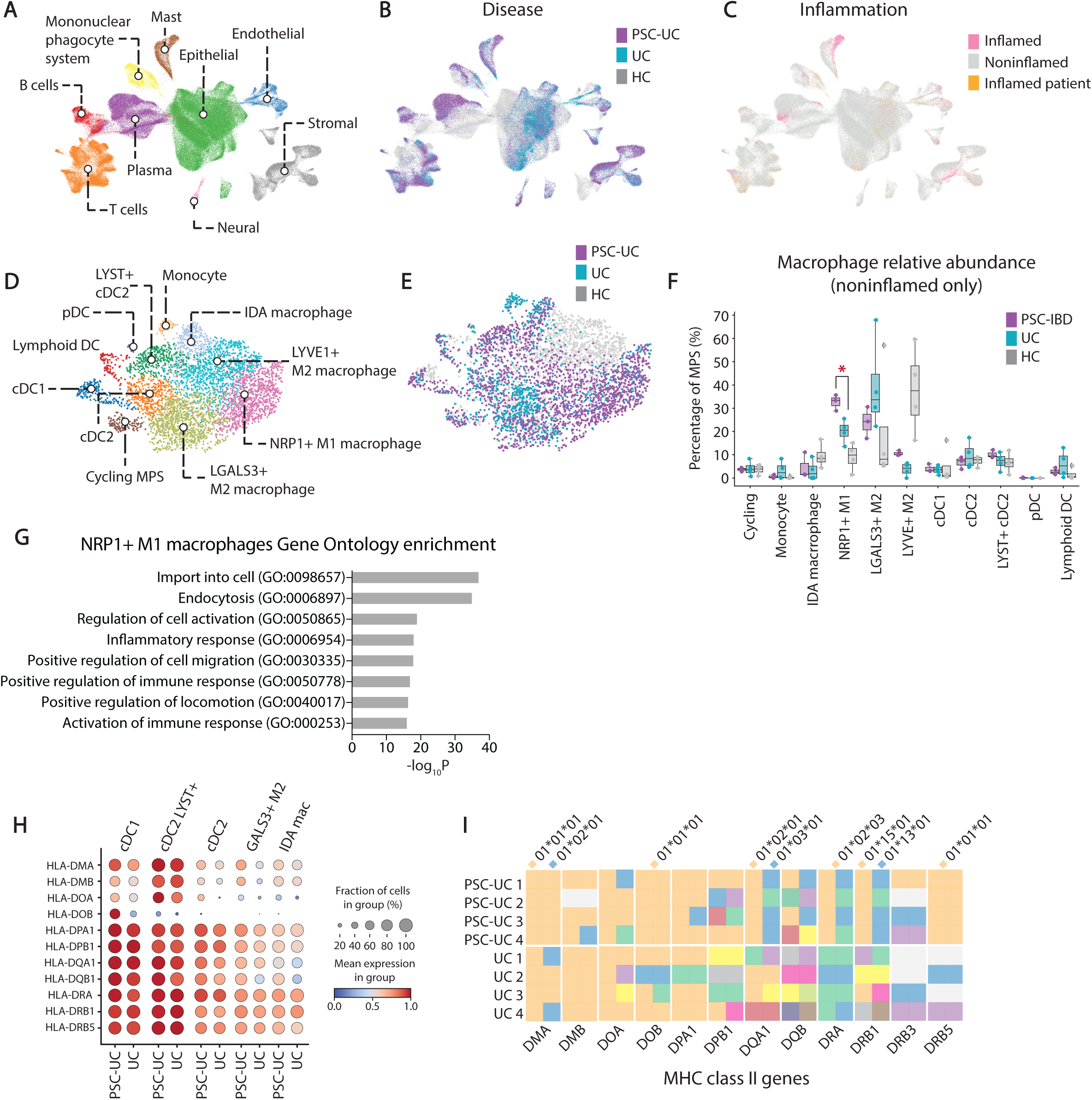
Conserved antigen presenting features of the PSC-UC gut. UMAP of scRNA-sequencing of colonic mucosa tissue from ascending, transverse, descending colon and rectum of healthy controls (HC), PSC-UC and UC patients (n=4/cohort) with clusters coloured by **A** | celltype lineage, **B |** cohort and **C |** inflammatory status of tissue. **D |** UMAP of the subsetted mononuclear phagocyte system (MPS) coloured by cell type annotation. DC, dendritic cell; pDC, plasmacytoid dendritic cell; cDC, central dendritic cell. **E |** UMAP of the MPS coloured by disease. **F |** Relative proportions of macrophage subtypes within the MPS compartment for PSC-UC, UC and HC. Each dot represents one patient (pooled 4 regional samples). *P* = 0.014 for M1 macrophage PSC-UC vs. UC, determined by unpaired T test. **G |** Enriched GO terms based on top 1000 upregulated genes for *NRP1+* M1 macrophages vs. rest of MPS. **H |** HLA class II gene expression across members of the MPS. **I |** Graphic visualisation of MHC class II gene allele expression from **Supplementary Table 3**. Colour denotes different alleles with key alleles annotated.

To explore the cellular diversity and function of the mononuclear phagocyte system (MPS) in the PSC-UC gut, we performed additional subclustering of the MPS compartment. We identified monocytes, three distinct conventional dendritic cell (cDC) populations, lymphoid DCs, plasmacytoid dendritic cells (pDCs), *NRG1*+ inflammation-dependent alternative (IDA) macrophages^15^, cycling myeloid cells, and three resident (*SELENOP*+, *C1QA*+) macrophage populations including two anti-inflammatory (M2-like) populations (*LGALS3+* M2 and *MRC1+* M2) and one *NRP1+* pro-inflammatory (M1-like) population (**Figure 2D-E**, **Supplementary Figure 2A** and **Supplementary Figure 3A**).

To assess differences in underlying disease biology, we first compared relative cell type proportions of biopsies taken from regions with no histological evidence of inflammation. Interestingly, PSC-UC patients displayed a significant enrichment of the *NRP1*+ macrophage population compared to UC and healthy cohorts (**Figure 2F**). This population was characterised by elevated expression of *SASH1*, pro-tumourigenic macrophage transcript *NRP1*^16^, and pro-inflammatory M1 markers *FOLR2* and *CD163L1*^15^ (**Supplementary Figure 3A**), and was enriched for inflammatory gene ontology terms (**Figure 2G**). *NRP1*+ macrophages also expressed *OTULINL* and *KLF2*, matching a population recently reported in the PSC liver^17^ (**Supplementary Table 2**, **Supplementary Figure 3A**). Together this indicates a disease-specific role for inflammatory macrophages in the PSC-UC gut, even during symptomatic remission.

Differential expression analysis revealed increased MHC class II transcript expression in dendritic cell and macrophage subtypes of PSC-UC patients compared to UC patients (**Figure 2H**). Given the previously reported association between PSC and specific HLA class II alleles^18^, we inferred the HLA genotypes of diseased donors from their single-cell expression data^19^. While our Australian PSC-UC cohort with caucasian ancestry did not express the ancestral AH8.1 haplotype as has been previously associated with PSC^20^, three out of four individuals expressed PSC risk alleles^20^ HLA-DRB1*13:01:01 and DQA1*01:03:01. Strikingly, all PSC-UC patients shared the DRB1*15:01:01 allele, DRA*01*02*03 allele, and were homozygous DMA*01:01:01, DOB*01:01:01 and DRB5*01:01:01 (**Figure 2I** and **Supplementary Table 3**). In contrast, UC patients displayed more variation in HLA class II allele expression (**Figure 2I**). Albeit in a small cohort of individuals, these results highlight a potentially stronger association between HLA class II allele expression and PSC incidence compared to UC.

Collectively, the enrichment of macrophages with a pro-inflammatory signature, upregulation of HLA class II transcripts in the MPS and sharing of HLA class II alleles in PSC-UC support a model whereby colitis in PSC is driven by an enhanced antigen-mediated inflammatory response.

### Lymphocyte patterning of the PSC-UC gut

Shared HLA class II allele expression among PSC-UC individuals suggests the involvement of antigen presentation to lymphocytes. We therefore characterised lymphocytes within the single-cell dataset. The conventional and unconventional T cell compartments were populated with innate lymphoid cells (ILCs), natural killer (NK) cells, γδ T cells, mucosal-associated invariant T (MAIT) cells and subtypes of CD4 and CD8 T cells (**Figure 3A-B, Supplementary Figure 2B**, and **Supplementary Figure 3B**). CD8 T cells included *GZMK*+ CD8 memory T and *CD69+* CD8 activated T cell subtypes. CD4 T cells included *SELL+* CD4 central memory T, *SELL+* T regularly (Treg), T follicular helper (Tfh), T helper (Th) 1, Th17, *CD69+* CD4 activated T and *ITGAE+* CD4 T cells.

**Figure 3.**
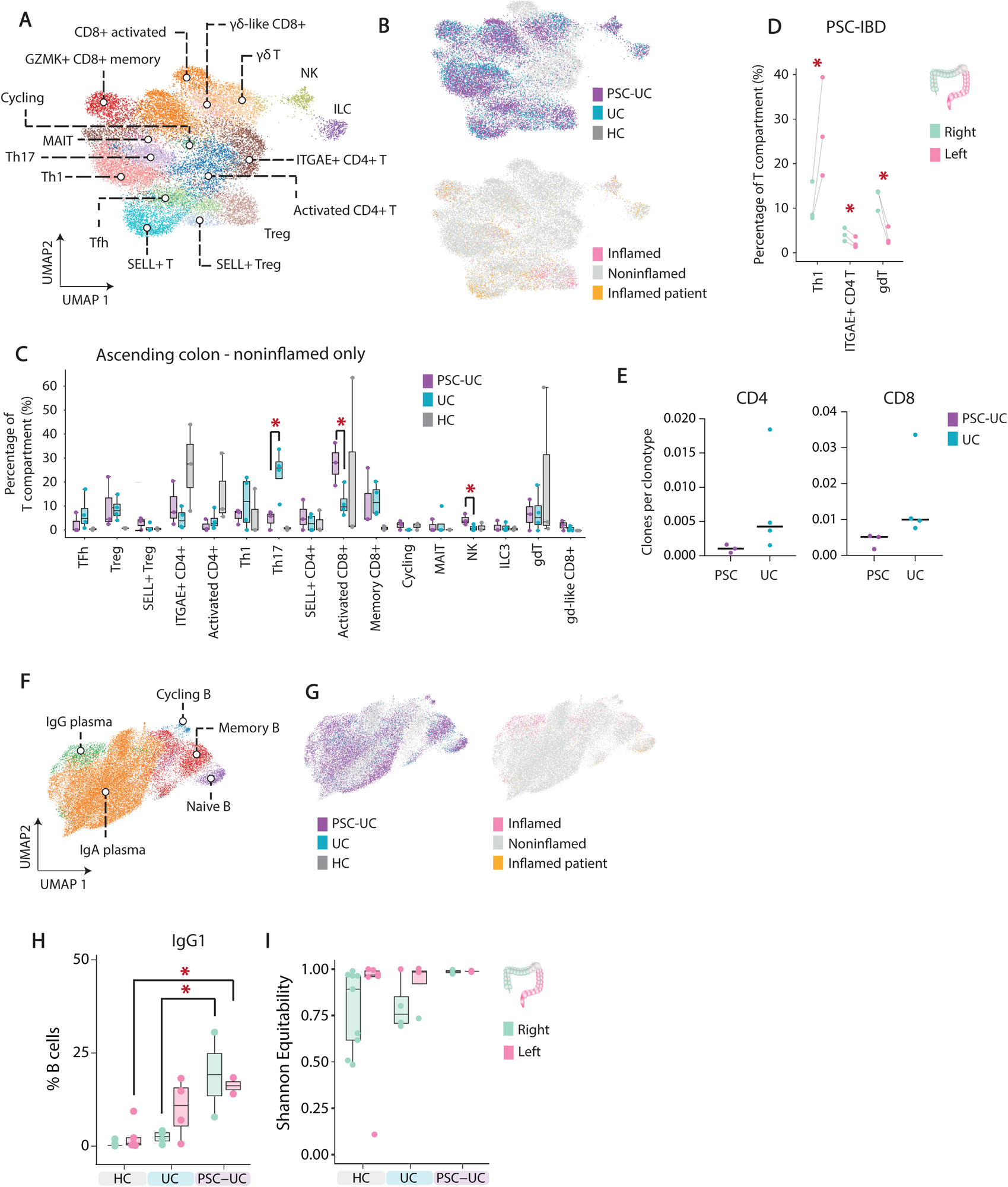
T and B cells in the PSC-UC gut. **A |** UMAP of the scRNA-sequencing dataset subsetted on T cell lineage and coloured by cell type. Th, T helper; Treg, T regulatory; ILC, innate lymphoid cell; NK, natural killer cell; MAIT, mucosal-associated invariant T cell. **B |** UMAP of the subsetting T lineage compartment coloured by disease and inflammation status. Inflamed patient refers to cells from patients experiencing regional inflammation that could not be accurately mapped to precise regions by hashtag demultiplexing. **C |** Relative abundance of each T cell subtype within the T cell lineage in the ascending colon (ACL) only. Each dot represents one patient biopsy. P values for PSC-UC and UC comparisons as determined by unpaired T test = 0.0226 for Th17, 0.0334 for activated CD8 T, 0.0484 for NK. **D |** Relative abundance of T cell subtypes in the left and right colon of PSC-UC patients. Each patient is joined by a grey line. Each dot represents pooled right (ascending and transverse) and left (descending and rectum) biopsies. P values as determined by paired T test = 0.05 (Th1), 0.021 (ITGAE+ CD4+ T) and 0.017 (gdT). **E |** Mean CD4+ and CD8+ T cell clone size in UC and PSC-UC patients. Each dot represents pooled data from one patient. **F |** UMAP of the scRNA-seq dataset subsetted on B cell and plasma compartment and coloured by cell type annotation. **G |** UMAP of the B cell and plasma compartment coloured by disease and inflammation status. **H |** Percentage of IgG1 B cells in the B cell compartment. P value via ANOVA + Tukey HSD = 0.015 for UC vs. PSC-UC right colons, 0.00018 for PSC-UC vs. healthy right colons and 0.04 for PSC-UC vs. control left colon. Each dot represents one patient. **I |** Shannon equitability of B cell clones. Each dot represents pooled cells of right (ascending and transverse) or left (descending and rectum) per patient.

To determine differences in underlying disease biology, we assessed relative proportions of T cell subtypes across each region of the colon in the noninflamed diseased and healthy data (**Figure 3C** and **Supplementary Figure 2C-E)**. This revealed an enrichment of activated CD8 T cells and NK cells and a lower proportion of Th17 cells in the ascending colon of PSC-UC compared to UC patients (**Figure 3C**). Additionally, when comparing right (pooled ascending and transverse) and left (pooled descending and rectum) colon, PSC-UC patients exhibited lower proportion of Th1 cells and increased levels of γδ T cells and *ITGAE*+ CD4 T cells in the right colon (**Figure 3D**). No significant differences in cell type proportions were detected between left and right colon in UC (**Supplementary Figure 2F**), which instead displayed an overall skewing towards Th17 cells relative to healthy controls as previously reported^21^. Matched TCR repertoire data identified a trend towards increased CD4 and CD8 T cell clonal expansion in UC compared to PSC-UC across both the left and right colon after normalising mean clonotype size to T cell numbers (**Figure 3E** and **Supplementary Figure 2G**). This supports that while the UC colon is characterised by a colon-wide clonally-expanded Th17 response, PSC-UC exhibits distinct anatomical patterning of unconventional lymphocytes and clonally diverse CD8 T cells, even in the absence of symptomatic inflammation. This patterning coincides with where disease pathology is most common in the PSC colon.

In the B and plasma cell compartments, we identified naive B, memory B, cycling B, IgA plasma and IgG plasma cells (**Figure 3F-G**, **Supplementary Figure 3C** and **Supplementary Figure 4A**). Analysis of matched B cell receptor (BCR) sequences showed that in the absence of active inflammation, isotype usage for both diseases was predominantly IgA1 and IgA2 (**Supplementary Figure 4B**), as seen in healthy control data and fitting with the mucosal nature of these samples^12^. Compared to healthy controls, both diseases had increased IgG1 usage at the left colon, however this was only significant for PSC-UC (**Figure 3H**). The most pronounced difference between diseases was at the right colon where on average 19.2% of PSC-UC B cells expressed IgG1 compared to just 2.7% for UC and 0.44% for healthy donors (**Figure 3H**). Curiously, while UC showed lower clonal antigen receptor diversity in the uninflamed right colon compared to the left colon consistent with healthy control, BCR diversity was equally high in the right and left colon of PSC-UC individuals without active inflammation, a signature predominantly driven by abundant IgA1/2 expressing cells (**Figure 3I, Supplementary Figure 4C**). Accordingly, more numerous expanded B clones (detected in >1 cell within a patient) were detected in UC, with a tendency to be within a single isotype compared to smaller expansions spanning multiple isotypes in PSC-UC (**Supplementary Figure 4D**). The greater clonal diversity and IgG1 preference of PSC-UC suggest a more acute B cell response particularly in the right colon compared to UC.

### A novel population of epithelial-endothelial intermediates in the UC gut

Given the growing appreciation for the role of stromal, neural and epithelial cell types in intestinal dysbiosis, we next compared for the first time at single-cell resolution these cellular compartments in the PSC-UC and UC gut. Within the epithelial compartment, we identified colonocytes, *AQP8+* absorptive colonocytes, transit amplifying cells, intestinal stem cells, goblet cells, *BEST4*+ epithelial cells and tuft cells (**Figure 4A-B, Supplementary Figure 3D** and **Supplementary Figure 4E-F**). Within the neural compartment, we identified glial cells, L cells, enterochromaffin cells and progenitors (**Figure 4C-D** and **Supplementary Figure 3E** and **Supplementary Figure 4G-H)**. Within the stromal compartment we identified myofibroblasts, *RSPO2+* myofibroblasts, pericytes, contractile pericytes, T reticular cells, and clusters corresponding to previously-reported populations Stromal 1, Stromal 2 and Transitional Stromal^11^. We also identified three further stromal populations, *CFH+* Stromal 1, *VSTM2A+* stromal and *CHI3L+* stromal cells, enriched in the PSC-UC and UC colon respectively (**Figure 4E-F, Supplementary Figure 3F** and **Supplementary Figure 4I**). The majority of cell states identified were at comparable frequencies between UC and PSC-UC. In line with increased fibrosis often observed in UC^22^, we observed a consistent enrichment in the proportion of myofibroblasts throughout the colon in UC versus PSC-UC when comparing patients in the absence of inflammation (**Figure 4G**).

**Figure 4.**
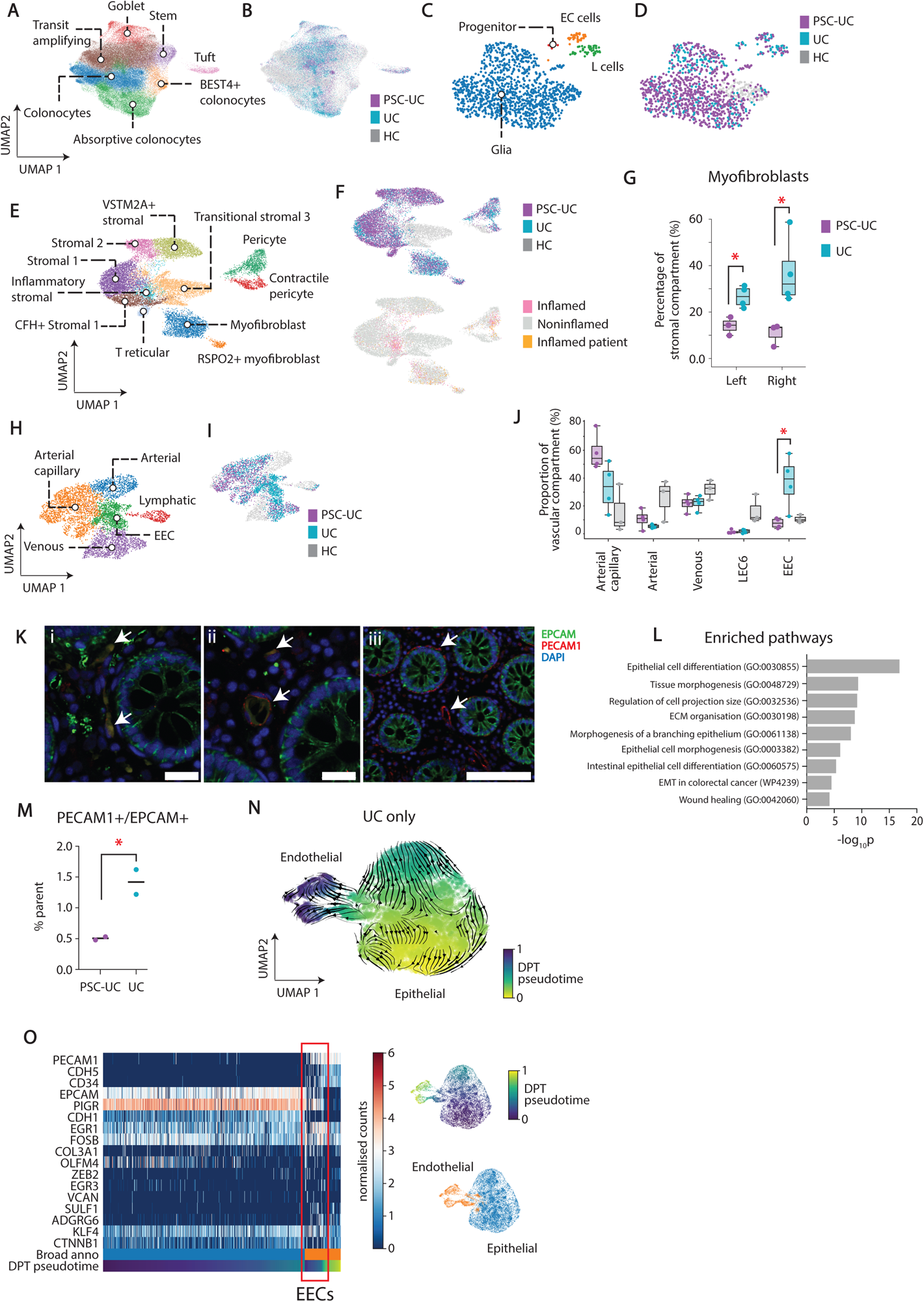
Stromal and endothelial cells of the UC gut. **A |** UMAP of the scRNA-sequencing dataset epithelial compartment coloured by cell type annotation and **B |** disease status. **C |** UMAP of the scRNA sequencing dataset neural compartment coloured by cell type annotation and D disease. **E |** UMAP of the scRNA-sequencing dataset stromal compartment coloured by cell type annotation. **F |** UMAP of the stromal compartment coloured by disease and inflammation status. Inflamed patient refers to cells from patients experiencing regional inflammation that could not be accurately mapped to precise regions by hashtag demultiplexing. **G |** Percentage of myofibroblasts in the stromal cell cluster of scRNA-sequencing of the right (pooled ascending and transverse colon) and left (pooled descending colon and rectum) in PSC-UC and UC. Each dot represents one patient. P values as determined by unpaired T test = 0.013 for left, 0.035 for right. H | UMAP of the scRNA-sequencing dataset endothelial compartment coloured by cell type annotation and **I |** disease status. **J |** Percentage abundances of cell types within the endothelial compartment. P value for PSC-UC compared to UC as determined by unpaired T test for epithelial-endothelial cells (EECs) = 0.023. **K |** Representative images from immunofluorescence imaging of colonic biopsy of a UC patient staining for DAPI (blue), EPCAM (green) and PECAM1 (red). Scale bars 20 µM for left and middle, 100 µM for right. **i |** Individual double positive cells. **ii |** Double positive cells arranged in channel-like structure. **iii |** Single-positive PECAM1 endothelial channel shows consistent extracellular PECAM1 staining and no EPCAM staining. **L |** Enriched pathway analysis terms for the top 1000 differentially-expressed genes of EECs relative to the endothelial cells. **M |** Percentage of double-positive EPCAM+/PECAM1+ cells among single, live cells in two UC and PSC-UC patients determined by flow cytometry. P = 0.0453 as determined by unpaired T test. **N |** UMAP showing epithelial (right cluster) and endothelial (left cluster) cells of UC patients, coloured by DPT pseudotime. Arrows depict splice variant velocity analysis. **O |** Expression of endothelial and epithelial markers as well as key genes of interest in cells shown in F ordered by pseudotime.

Within the vascular compartment, we identified pericytes, contractile pericytes, capillary, arterial, venous and lymphatic endothelial cells (**Figure 4H, Supplementary Figure 4J-K** and **Supplementary Figure 3G**). We also detected an unexpected population of endothelial cells with high expression of canonical epithelial markers (*EPCAM+, PIGR+*), that were significantly enriched in UC compared to PSC-UC and healthy control biopsies, hereafter termed epithelial-endothelial cells (EECs), (**Figure 4I-J**, **Supplementary Figure 5A**). The cluster was stable following the implementation of stringent doublet removal algorithms and did not display increased gene or count numbers (**Supplementary Figure 5B**). Immunofluorescence imaging (**Figure 4K** and **Supplementary Figure 5C**) of UC colonic biopsies confirmed the existence of individual EPCAM+/PECAM1+ cells outside of villi structures (**Figure 4Ki**) and organised into channel-like structures resembling blood vessels (**Figure 4Kii**). Within the same patient, typical single-positive PECAM+ vessels were also detected (**Figure 4Kiii**). Intriguingly, EPCAM staining appeared to localise to the cytosol in EPCAM+/PECAM1+ cells, fitting with an epithelial-mesenchymal transition (EMT)-like program where membrane-bound EPCAM is cleaved and its intracellular domain translocated to the nucleus to drive EMT-related gene expression^23^. Accordingly, gene ontology analysis of differentially expressed genes of the double-positive population compared to other endothelial cells identified an enrichment of pathways corresponding to differentiation, morphogenesis, wound healing and EMT (**Figure 4L**, **Supplementary Table 4-5**). EECs were not detected by immunofluorescence in healthy tissue (**Supplementary Figure 5C**). We next quantified the proportion of membrane EPCAM+/PECAM1+ cells in PSC-UC and UC patients by flow cytometry, supporting a consistent enrichment of EPCAM+/PECAM1+ cells in the UC gut (1.42% versus 0.5%, mean from two independent experiments, **Figure 4M, Supplementary Figure 5D**).

Previous work has reported dysregulated vasculature in the UC gut^24,25^. Therefore, we hypothesised that an epithelial-to-endothelial transition (EET) may contribute to vascular dysregulation in UC, similar to vasculogenic mimicry programmes observed in cancer^26^. To explore this, we subsetted the UC epithelial and endothelial compartments, excluding goblet cells, tuft cells, BEST4+ colonocytes and lymphatic endothelial cells, for further analysis of their cellular trajectories (**Supplementary Figure 6A**). Diffusion pseudotime and splice variant velocity analysis implicated EPCAM+/PECAM1+ cells in a differentiation trajectory between the epithelial and endothelial state (**Figure 4N**). Ordered by pseudotime, EECs display a gradual downregulation of epithelial markers *EPCAM* and PIGR alongside a concurrent upregulation of endothelial markers *PECAM1* and *CD34* (**Figure 4O**). Notably, *EPCAM* downregulation appears to occur before *PIGR* downregulation, suggesting that cleavage of EPCAM is an initiating factor in this transition. While the expression of most major EMT transcription factors were not enriched in EECs (**Supplementary Figure 6B**), *ZEB2* expression was detected (**Figure 4O**). In addition, EECs upregulated the intestinal stem marker *OLFM4*, previously identified as upregulated in the UC gut^15^, and *EGR1* (**Figure 4O** and **Supplementary Figure 6A**), reported to drive vasculogenic mimicry and angiogenesis in various cancers^27,28,29^ and the differentiation of mesenchymal stem cells into endothelial cells^30^.

We next queried what cells might be signalling to EECs and contributing to their development. Analysis of ligand-receptor expression with CellphoneDB revealed EEC expression of *ADGRG6* (**Supplementary Figure 6C**), an adhesion G protein coupled receptor most studied in the context of Schwann cell maturation. Upregulation of *ADGRG6* has recently been shown to promote breast^31^, bladder^32^ and colon^33^ cancer cell growth and angiogenesis^34^ as well as developmental and pathological angiogenesis in the eye via STAT5 and GATA2-mediated control of *VEGFR2* expression^35^. Predicted *ADGRG6* ligands include *COL4A6* and *COL4A3*, expressed by fibroblasts, and *LAMA2*, expressed by myofibroblasts, which were particularly enriched throughout the UC colon (**Figure 4A**). EECs were also enriched for the integrin α6b4 complex (*ITGA6* and *ITGB4* transcripts). ITGA6 is a marker of tumour endothelial cells in hepatocellular carcinoma^36^, and the integrin α6b4 complex has been shown to induce *VEGF* and FGF-mediated tumour angiogenesis^37,38^. Neuregulin-1 (*NRG*), highly expressed by fibroblasts in our dataset, is a ligand for the integrin α6b4 complex. Together, the identification of UC-specific EECs points toward an epithelial-to-endothelial tissue remodelling programme occurring in the gut of UC patients.

### PSC-UC and UC intestinal inflammation is characterised by TMEM176B+ mast cell infiltration

We next assessed PSC-UC and UC in the context of inflammation. Active inflammation in both diseases was characterised by an enrichment of Tregs, TFhs and *CHI3L1*+ inflammatory fibroblasts (**Supplementary Figure 7A-B**). In noninflamed individuals, we identified a population of mucosa resident mast cells and a small population of *CLC+* mast cells (**Supplementary Figure 7C**). In inflamed patients, we also identified a transcriptionally distinct mast cell state, hereafter termed inflammatory mast cells (IMCs, **Figure 5A**) that was only present in the colons of PSC-UC and UC patients experiencing inflammatory relapse compared to remission (containing ‘resident’ mast cells only). Both mast cell subsets expressed tryptases (*TPSAB1* and *TPSB2*), carboxypeptidase (*CPA3*) and low levels of chymase (*CMA1*) (**Figure 5C**), in contrast to the classical ‘mucosal’ signature of tryptase only^39,40^. Interestingly, IMCs displayed significantly decreased expression of the relatively unstudied delta tryptase enzyme compared to resident mast cells (**Figure 5C**). Furthermore, both subsets expressed the FcERI receptor, but no other immunoglobulin-activated receptors (**Figure 5C**). Pathway analysis of IMC versus resident mast cell differentially-expressed genes (**Figure 5D**) identified enrichment of terms involved in leukocyte activation, degranulation, vesicle-mediated transport and FC epsilon receptor signalling (**Figure 5E**), together pointing to an active phenotype in IMCs. We did not detect IgE transcript expression in plasma cells of inflamed patients, though we did detect small amounts in B cells mostly in UC patients (**Supplementary Figure 6D**). However, in the single cell V(D)J sequencing data, no B cells or plasma cells were observed to express IgE immunoglobulin heavy chain transcripts.

**Figure 5.**
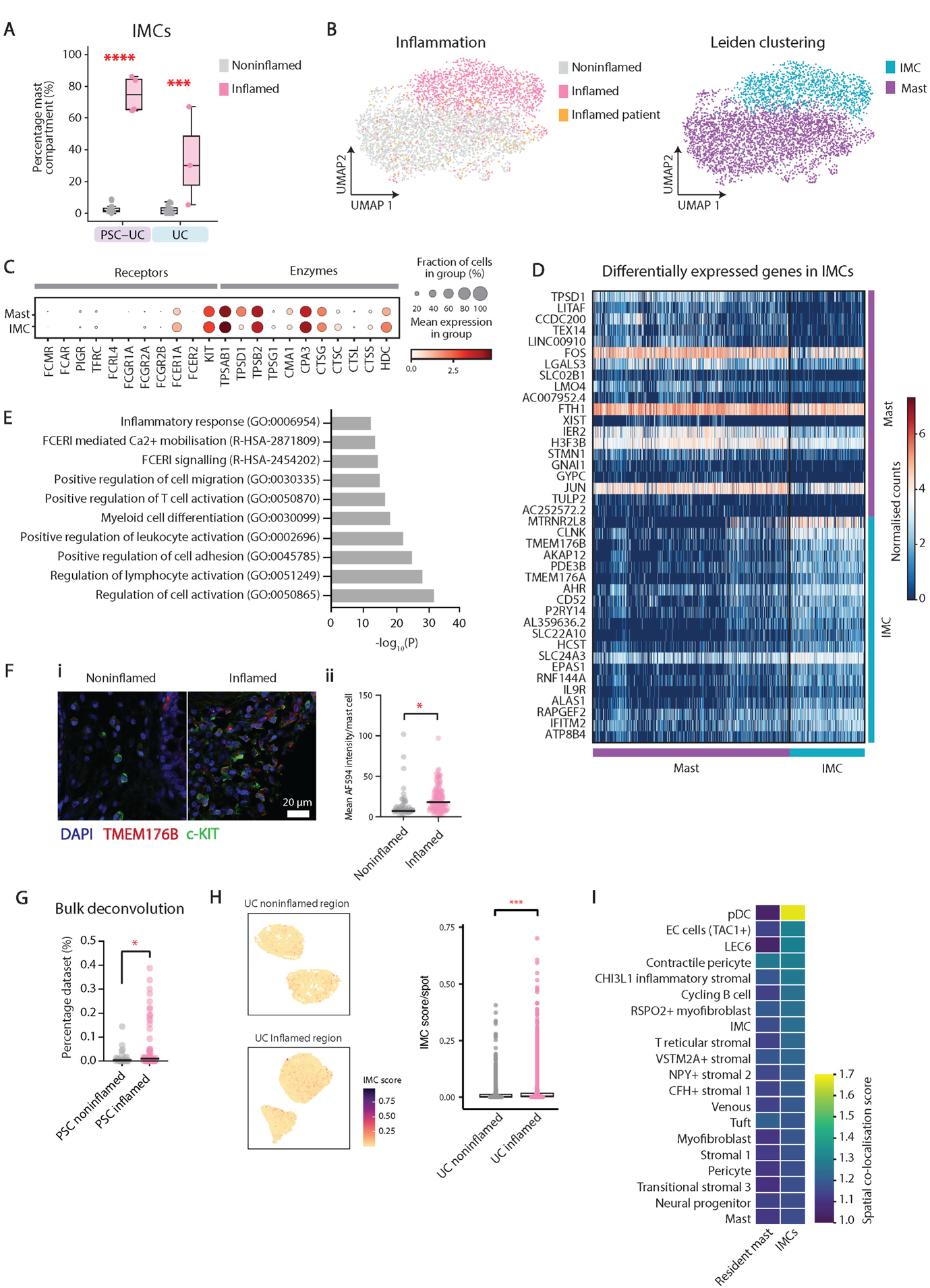
UC and PSC-UC inflammation is characterised by infiltration of *TMEM176B+* mast cells. **A |** Percentage of inflammatory mast cells (IMCs) in the mast cell compartment of the scRNA-sequencing dataset for PSC-UC and UC patients. Each dot represents one biopsy. P values = 1.25×10^-13^ for PSC-UC and 0.00023 for UC as determined by unpaired T test. **B |** UMAP of mast cells and IMCs coloured by tissue inflammation state and leiden clustering. ‘Inflamed patient’ refers to cells from patients experiencing inflammation in one or more regions, but the localisation (and therefore inflammatory state) of the individual cell could not be accurately determined by hashtag demultiplexing. **C |** Dot plot depicting expression of common mast cell receptors and enzymes in mast cells and IMCs. **D |** Heatmap displaying cellular normalised expression values of the top 25 up and down regulated genes in IMCs vs. mast cells. **E |** Enriched pathway terms in IMCs based on the top 1000 upregulated genes in IMCs vs. mast cells. **F | i** Representative Immunofluorescence images of noninflamed and inflamed PSC-UC tissue. **ii** Mean AF594 fluorescence intensity (TMEM176B staining) per mast cell defined by DAPI and c-KIT using Imaris software. P value as determined by unpaired T test = 0.017. **G |** Spatial sequencing of inflamed and noninflamed UC tissue deconvoluted using our scRNA-seq dataset and coloured by IMC score (left) and quantified (right). **H |** Co-localisation scores generated from the inflamed UC spatial transcriptomics dataset for resident and inflammatory mast cells (IMCs). Top 20 highest scoring cell types co-localising with IMCs. P value as determined by Wilcoxon Rank Sum = 3.56×10^-31^ **I |** Deconvolution of Shaw *et al*.’s^9^ PSC-IBD bulk RNA sequencing dataset using our own scRNA-seq data identified an enrichment of an IMC signature in PSC-IBD patients with intestinal inflammation. P value as determined by unpaired T test = 0.0028.

Compared to resident mast cells, the IMC gene signature was characterised by expression of numerous transcripts associated with FcERI-mediated activation, such as cytokine-dependent hematopoietic cell linker (*CLNK*)^41^, hematopoietic cell signal transducer (*HCST*, encoding the DAP10 protein) and PDE3B^42^ (**Figure 5E** and **Supplementary Table 6-7**). However, for each of these transcripts, genes encoding known interacting partners and downstream mediators (PLCG1, GRB1, LAT for CLNK^41^ and KLRK1, SIRPB1 for HCST^43,44,45^) were not enriched in IMCs, pointing either toward a yet uncharacterised activation pathway occuring during intestinal inflammation or discrepancies between mRNA expression and protein levels. More recently, *HCST* expression has been described in tumour-infiltrating mast cells in renal cell carcinoma^46^. IMCs were also characterised by high expression of ion channel transcripts *TMEM176A* and *TMEM176B,* the latter validated at the protein level in PSC-UC inflammation by immunofluorescence (**Figure 5F** and **Supplementary Figure 7E**). The IMC signature was also observed in an independent inflamed UC single cell RNA seq study^15^ (**Supplementary Figure 7F**) and enriched in deconvoluted inflamed versus noninflamed PSC-IBD bulk RNAseq data (**Figure 5G**)^9^.

Spatial transcriptomic analysis of inflamed versus noninflamed UC colon biopsies further supported the enrichment of IMCs in inflammation (**Figure 5H**). Furthermore, deconvolution of all cell type signatures from our single cell dataset showed IMCs most commonly colocalising with pDCs and enterochromaffin cells (**Figure 5I**) and with venous, pericyte and LEC6 endothelial cells fitting with their positioning near vessels^40^. Together, this suggests IMCs support an active immune cell environment in the inflamed IBD intestines.

### TMEM176B is involved in mast cell degranulation

To assess whether an IMC signature could be recapitulated *in vitro*, we performed qPCR to measure the expression of activation (*Tnf, Il6*) and IMC (*Tmem176b*, *Clnk*) markers on the Cl.MC/C57.1 mast cell line^47^ at various time points following FcERI-mediated activation (**Figure 6A-B**). This revealed a reduction in both *Tmem176b* and *Clnk* following stimulation with cognate antigen compared to negative controls, suggesting that IMCs may represent an influxing population of yet unstimulated mast cells. We next assessed the role of TMEM176B in mast cell activation via the use of pharmacological inhibitor BayK8644^48^. Inhibition of TMEM176B resulted in decreased degranulation following FcERI-mediated stimulation, suggesting that TMEM176B may play a functional role in mast cell activation during intestinal inflammation (**Figure 6C**). Furthermore, the IMC and resting mast cell signatures scored highly as contributing to a PSC-IBD-specific inflammatory signature^9^ predictive of the likelihood of developing dysplasia (**Figure 6D**). Lastly, mast cells expressing *TMEM176B* (as well as IMC-specific transcript CD52) have been shown to be enriched in colorectal cancer^49^, and *TMEM176B* expression correlates with decreased survival in this patient group (**Figure 6E**). These analyses point toward a pathological role for IMCs in colon tumourigenesis.

**Figure 6.**
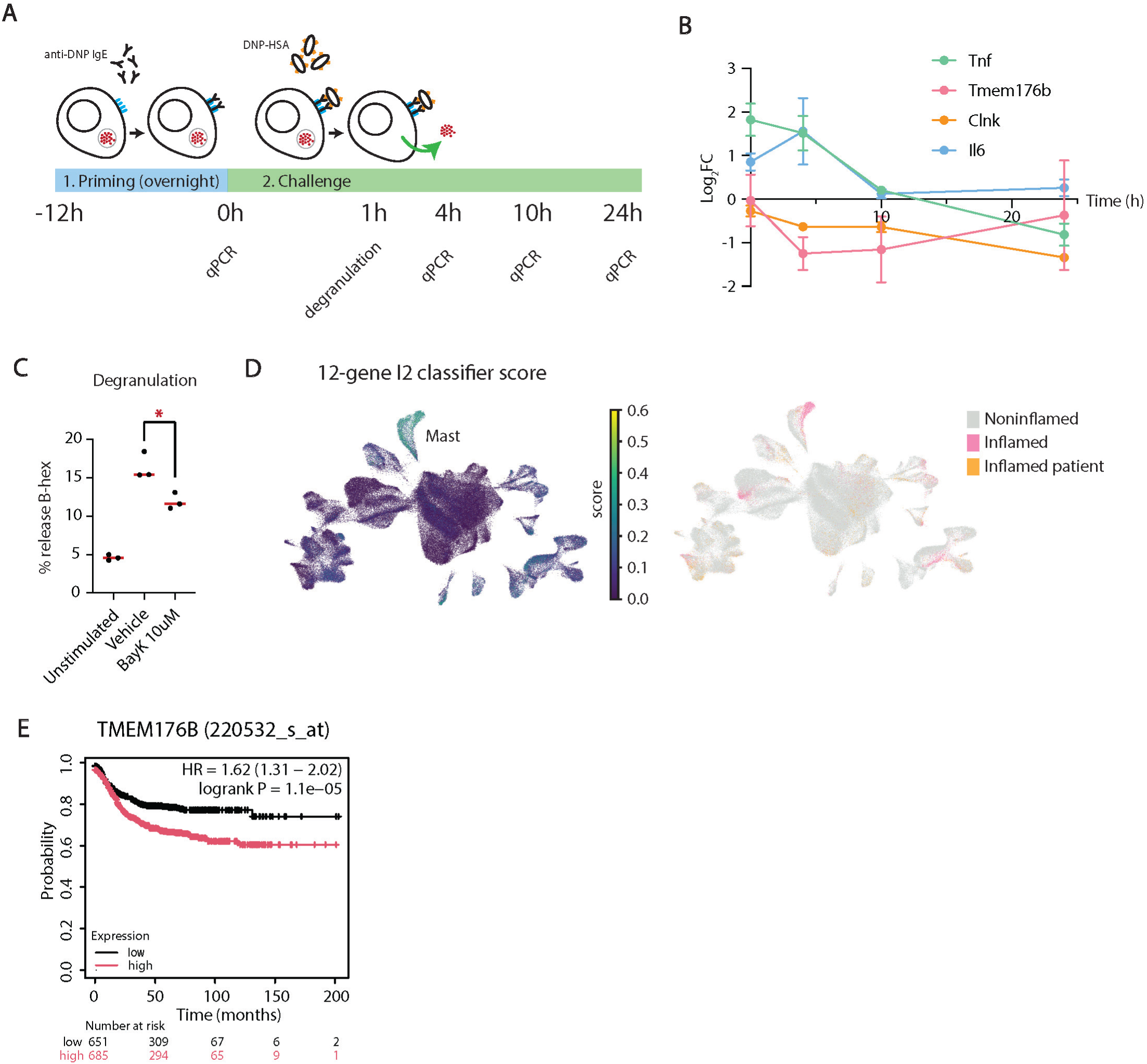
TMEM176B is a marker of sustained activation. **A |** Experimental design to assess gene expression by qPCR and degranulation by quantification of β-hex in supernatant. **B |** Time course expression of *Tnf*, *Tmem176b*, *Clnk* and *Il6* in murine mast cells following activation via FCERI stimulation. **C |** Mast cell degranulation following FCERI-mediated activation in the presence of vehicle and TMEM176B inhibitor BayK8644. P value as determined by independent T test = 0.0191. **D |** UMAPs coloured by expression of Shaw *et al*.’s 12-gene I2 classifier gene list predictive of CRC in PSC-UC (left) and inflammation (right). **E |** Kaplan-Meier plot for TMEM176B expression in colorectal cancer.

## Discussion

A challenge in developing targeted therapeutics for PSC-UC is a limited understanding of how the pathogenesis compares to that of IBD. Previous transcriptomic studies have shown differences at cellular and microbial levels between PSC-IBD and UC, but with limited appreciation of the contribution of diverse cell lineages to intestinal immunity^10^ and consideration of distinct anatomical locations. To this end, we performed the first scRNA-seq study on unsorted cells of mucosal biopsies of PSC-UC and UC colon, coupled with matched 16S amplicon sequencing, spatial transcriptomics and *in vitro* validation. Our analysis spans four anatomical colon regions and active and remittent inflammation, providing a platform for multi-cell lineage comparisons across IBD subtypes, anatomical space and disease states. Importantly, we show for the first time that the cellular landscape of the PSC-UC and UC colon are inherently different in the absence of active disease.

By analysing the mucosal-adherent microbiome of four anatomical regions, we have shown that PSC-UC patients exhibit reduced diversity compared to UC patients specifically in the ascending colon, the region in which both inflammation and neoplasia most commonly develop^50^. Reduced diversity in the ascending colon could be due to the downstream effects of defective bile acid production and lipid metabolism^51^ in the small intestine of PSC-UC patients. This finding contrasts with previous studies reporting no difference in mucosal microbial diversity between UC and PSC-IBD at different colon locations^14,52,53^. These discrepancies may be due to differences in inflammatory status of biopsies across the studies. Previous studies have similarly reported enrichment in *Pseudomonas*^6^ and *Blautia*^52^ in the PSC-UC mucosal microbiome. Limited overlap in bacterial populations between our study and previous works may be due to the effect of geographical location-no previous studies have characterised the Australian PSC-UC mucosal microbiome. Our observation of a more conserved microbiome in UC compared to PSC-UC suggests that in PSC-UC, the microbiome composition may play a less dominant role in intestinal dysbiosis than in UC. Nevertheless, further work is needed to decipher whether the microbial alterations described in PSC-UC are an initiating factor for colitis in PSC patients, or concomitant with cholestatic liver disease in general.

We determined marked sharing of HLA class II alleles by PSC-UC patients, as previously reported^20,54^, which was not evident in the UC patients. While this is indicative of an autoimmune response and indeed PSC and UC are considered atypical autoimmune diseases, our TCR and BCR sequencing analyses did not reveal significant clonal expansions. Future analysis of liver biopsies from PSC patients could shed light on whether an autoimmune response occurs in the liver of PSC patients prior to the onset of gut disease. Our analysis of the T cell compartment supported a contribution of Th17 cells throughout the colon of both diseases^6,55^. However, uniquely in our study, we revealed a distinct cell signature in the right colon of PSC-UC patients marked by increased proportions of NK, *CD8+* T, *ITGAE*+ Th1, and γδ T cells. Previous works have identified expanded γδ T cells in the peripheral blood^56^ and liver^57^ and NKT-like cells in the liver^17^ of PSC patients, pointing toward shared cellular mechanisms between liver and gut disease manifestations. Furthermore, we identified a PSC-UC enriched population of *NRP1*+ tissue-resident macrophages that shares some markers (*OTULINL*, *KLF2*) described in the recently-reported PSC-enriched liver macrophage population^17^. While tissue-resident macrophages are not thought to migrate between organs, gut and liver *NRP1+* macrophages may share a common developmental origin, providing another link between gut and liver inflammation in PSC. From these findings it is tempting to speculate that a yet unknown autoantigen initiates inflammation of the biliary tree and disrupts bile acid secretion, which results in an altered microbiome^58^ most notably in the proximal colon. This in turn drives a distinct local immune cell response which contributes to the right-sided dominance of colon inflammation and perpetuates liver fibrosis.

Our unbiased single-cell approach and sampling of noninflamed tissues allowed for the detection of a rare, hybrid epithelial-endothelial cell state. This cell state may have been missed from previous scRNA-seq datasets profiling the inflamed UC gut^15^ where samples are enriched for immune cells. Accordingly, we did not detect EECs in samples from inflamed regions. The notion that intestinal epithelial cells may transdifferentiate to endothelial-like cells in UC supports the ‘leaky gut’ theory of intestinal inflammation^59^, whereby increased intestinal permeability and exposure of pro-inflammatory microbial and dietary compounds to circulating immune cells drive inflammation. Interestingly, despite IBD patients exhibiting increased and dysregulated vasculature^25,60^, many drugs inhibiting the classical VEGF/VEGFR angiogenesis pathways have been shown to exacerbate IBD^61–63^, while the activation of the classical VEGF-C/VEGFR3 pathway is protective^64^. In another study, where the use of antiangiogenic receptor tyrosine kinase inhibitors effectively suppressed the VEGF/VEGFR pathway, no decrease in colonic microvessel density was observed^65^, pointing toward the existence of a non-canonical vasculogenic programme. An epithelial-to-endothelial transition, which becomes the dominant vasculogenic programme following VEGF/VEGFR inhibition, could explain the poor therapeutic success of classical anti-angiogenic drugs and the comparative success of anti-TNF therapies (which inhibit EMT and cellular plasticity), as well as the observation of immature, leaky vessels with little pericyte coverage in IBD patients^25^.

We identified a conserved *TMEM176B+* activated inflammatory mast cell subtype in the current and previously-published IBD datasets. Interestingly, we detected more *IGHE* expression in B cells from UC patients, who exhibited fewer IMCs. This may be due to differences in the time of sampling, as IgE-expressing B and plasma cells are rare and mostly short-lived^66,67^. Due to the sterile transcription of the *IGHC* genes, detection of *IGHC* transcripts (such as IGHE) does not perfectly correlate with the presence of B cell isotype expression^68^. Increased expression of the *IGHE* transcript could be indicative of a higher frequency of IgE+ B cells, or could suggest that they may be poised for potential switching to IgE as transcription keeps the locus accessible for class-switch recombination^69^. Furthermore, previous works have demonstrated elevated IgE in the blood of PSC patients^70–72^, and it is possible that IgE generated outside the gut may have generated an IMC response in PSC-UC, in-line with the trend of different underlying triggers of PSC-UC and UC converging to similar inflammatory mechanisms. The detection of IgE-activated mast cells during IBD flare raises interesting questions regarding the role of mast cells in autoimmunity. Indeed, recent studies have demonstrated IgE involvement in non-allergic autoimmune disease, as well as in tumour progression^73^. The enrichment of TMEM176B+ mast cells in PSC-UC, TMEM176B expression by CRC mast cells, and a likely contribution of mast cells to a pro-dysplasia inflammatory signature in PSC-IBD provide compelling evidence for the involvement of mast cells in colon tumourigenesis, and may constitute a valuable therapeutic or predictive target.

Analysis of patient samples in this study presents notable limitations. Firstly, IBD patients commonly have diverse therapeutic histories which can impact cellular responses and microbiome composition, though we did not include patients with antibiotic treatment in the 4 weeks prior to sampling. Bowel preparation prior to colonoscopy is also likely to reduce microbial diversity. Additionally, while PSC is usually diagnosed prior to the onset of colitis^5^, it is possible that UC patients included in this study go on to develop PSC. Such cases would shed more light on whether the distinct right colon cell profile of PSC-UC is the result of or contributing to liver disease. We used the best available murine mast cell line that expresses high FcεRI and key markers such as KIT (CD117), CD33 and CD203c^74^ as isolating and culturing patient-derived primary mast cells was not feasible and none of few human mast cell lines available to date^75^ were suitable for this study. Future work may look to animal models to further study the role of IMCs in intestinal inflammation, though these present their own challenges in their relatability to human intestinal disease. Finally, while our sample size is relatively small (n=4/disease), particularly for PSC patients with inflammation, our study still represents the most comprehensive single cell analysis of the PSC-UC gut to-date.

## Conclusion

We have generated the first unbiased parallel single cell mRNA and bacteria sequencing dataset of the PSC-UC and UC large intestine. In doing so, we have demonstrated that the PSC-UC and UC colons are microbially and cellularly distinct even in the absence of active disease. In particular, PSC-UC patients display distinct anatomical patterning not seen in UC, with reduced microbial diversity and unique immune cell enrichments in the right colon. We describe a rare, epithelial-endothelial hybrid cell population which may contribute to intestinal permeability in UC, further exemplifying the cellular distinctness of the UC and PSC-UC gut. Finally, we identify a shared *TMEM176B+* activated mast cell response during inflammation and provide evidence for these cells in intestinal tumourigenesis. The disease-specific cell and microbial signatures identified from this work represent tools for patient stratification and avenues for precision therapies.

## Methods

### Sample collection and patient cohort

Mucosal pinch biopsies were collected from the ascending colon, transverse colon, descending colon and rectum of patients with either UC or PSC-UC (classified as resembling UC at the clinical level) during routine endoscopy procedures. Prior to endoscopy, *Picoprep* bowel preparation was used, which contains active ingredients sodium picosulfate, magnesium carbonate hydrate, and citric acid. All patients were Australian and of typically white European descent and had no antibiotic use in the 28 days prior to endoscopy. Inflammatory state was recorded by treating gastroenterologists during endoscopy and histologically confirmed by anatomical pathologists with subspecialty interest in gastrointestinal pathology. Mucosal pinch biopsies were immediately transferred to ice-cold RPMI medium containing 10% FBS. All patients were recruited as part of the Australian IBD Microbiome (AIM) study. All procedures were covered by ethics approval from South Eastern Sydney Local Health District Research Ethics Committee (HREC ref no:18/173) and followed patient consent.

### 16S rRNA gene sequencing

Mucosal pinch biopsies were collected from the ascending colon, transverse colon, descending colon and rectum. Biopsies were washed in 1x PBS and cryopreserved at -80C in FBS containing 10% DMSO. For DNA isolation, biopsies were washed in PBS and digested overnight in a thermoshaker at 700 rpm/56°C in 180 µL Buffer ATL (QIAGEN) with 30 µL proteinase K (QIAGEN). DNA extraction was then performed using the QIAamp DNA Mini Kit (QIAGEN) as per the manufacturer’s instructions. The V3-V4 hypervariable region of the 16S rRNA gene was targeted for amplicon sequencing using the 341f-805r primer pair as outlined in Illumina’s (San Diego, CA) 16s Metagenomic Sequencing Library Preparation guide. Due to low microbial biomass in biopsy samples, 5 PCR reactions per biopsy sample were run in the 1^st^ stage PCR. These 5 reactions were combined during the PCR clean-up stage. Second-stage PCR and subsequent PCR clean-up were undertaken on single reactions. Samples were sequenced on Illumina MiSeq at the Ramaciotti Centre for Genomics (UNSW Sydney), generating paired-end 300bp reads.

### Analysis of microbial amplicon sequencing

Amplicon data were quality controlled with dada2^76^ embedded in qiime2^77^. Host contamination was removed using Bowtie 2 (version 2.4.2)^78^. Taxonomic annotation was performed using the qiime2 feature classifier plugin with the Silva 138 database. R (v4.2.2) packages qiime2R (https://github.com/jbisanz/qiime2R) and phyloseq^79^ were used for diversity analyses. The Mann-Whitney-Wilcoxon and Kruskal-Wallis tests were applied in the comparisons of feature abundance and alpha diversities between groups. Adonis (from the Vegan package) based on Bray-Curtis and Jaccard distances was used to investigate and rank the effect of disease on overall microbial composition. P values were adjusted for multiple testing where appropriate using the Benjamini-Hochberg method. Relative abundance analysis was performed for each biopsy by calculating the number of counts for each bacterial genus divided by the total number of counts. Genus level counts were used to generate an Anndata object with metadata as observations and genera as variables. Anndata objects were used to run differential abundance analyses in SCANPY (version 1.9.1) after normalising counts per biopsy and log transformation.

### Single cell RNA-sequencing library preparation

For single cell RNA sequencing, 2-3 biopsies per region were immediately processed following isolation. All patients were processed individually, except for PSC-2 and PSC-3 who were processed in parallel. Each region was processed separately until pooled for capture. Biopsies were washed in 1x PBS before incubation in 1.5 mL digestion medium (1.16 mL HBSS, 246.9 µL liberase DH, 60 µL hyaluronidase, 36 µL DNase I) in 15 mL tubes with rotation at 600 rpm for 25 minutes at 37°C. Biopsies were homogenised at 10 minutes via trituration and vortexing. Digestions were halted by adding 5 mL ice-cold neutralisation medium (20% FBS in RPMI) and cell suspensions passed through 40 µM cell strainers. Digestion tubes were washed with a further 5 mL neutralisation medium passed through the same strainer to transfer any remaining cells. Cell suspensions were then centrifuged at 300g for 10 minutes at 4°C. Supernatants were discarded and cells resuspended in 1 mL resuspension medium (1% FBS in PBS) and centrifuged again as before. Cells of the right colon (ascending and transverse colon) and left colon (descending colon and rectum) were pooled separately, with hashtag oligos used for *in silico* deconvolution of discrete anatomical regions for each donor. Supernatants were then discarded and cell pellets resuspended in 45 µL Cell Staining Buffer with 5 µL TruStain Fc Blocking Reagent and incubated for 10 minutes at 4°C to prepare for hashing. TotalSeq C0251 anti-human hashtag 1 and C0252 anti-human hashtag 2 antibody pools were prepared as per the manufacturer’s instructions and added to 50 µL blocked cell suspensions and incubated for 30 minutes at 4°C. Hashed cell suspensions were then washed three times by centrifugation at 400 g at 4°C for 5 minutes with 3 mL Cell Staining Buffer. Cell pellets were then resuspended in 100 µL and counted. Regions were pooled equally for two captures: right colon (containing ascending and transverse colon) and left colon (containing descending and rectum). Demultiplexing based on antibody hashtagging was therefore used for higher-precision mapping to each region (e.g. descending colon vs. rectum). For PSC-2 and PSC-3, regions were pooled across 4 captures and cells demultiplexed using SNP information.

### Preprocessing of scRNA-seq data

The Cell Ranger (v7.0) analysis pipeline using *cellranger multi* with default settings was used to generate counts files from gene expression and V(D)J library files. Loom files containing spliced and unspliced gene expression matrices were generated using the command line interface tool Velocyto^80^ as per the standard usage guide. Splice variant data generated for each run was subsequently read into Anndata files for each run using the scv.utils.merge() function from the scVelo package (version 0.2.4)^81^. HTO demultiplexing for each run was performed in R and read into the Anndata object for each run. For patients PSC-2 and PSC-3 who were captured together, SNP information was used to demultiplex patients using the Souporcell tool^82^ within the Demuxafy framework^83^ and information read into the Anndata object. All further single cell analyses were performed using the Scanpy package^84^ (v1.9.1) in Python (v3.9.4) with all function parameters at default settings unless described otherwise. Filtered counts files from each capture were first subjected to doublet detection using Scrublet with a manually-adjusted doublet score cut-off threshold determined using the scrub.plot_histogram() function. Individual counts matrices were merged into a single AnnData object and cells were filtered for more than 200 genes and less than 50% mitochondrial reads. Genes expressed in less than three cells were also removed, as well as mitochondrial and ribosomal genes. At this point, cells from inflamed regions were either removed or retained, depending on the analysis. The dataset was then normalised and log transformed before being filtered for highly variable genes determined using the standard Scanpy workflow. The number of counts per cell and percentage mitochondrial gene expressed were regressed before data scaling and principal component analysis. Batch-corrected nearest neighbours and dimensionality reduction were performed using Scanpy’s bbknn and UMAP functions.

### Cell type annotation of scRNA-seq data

Leiden clustering was first performed at resolution 0.3 on full datasets and highly expressed genes for each cluster were defined using the sc.tl.rank_gene_groups() function and used to manually annotate broad cell lineages based on established marker genes from previously published data. Broad cell lineage annotations were then read into the original, non-normalised counts files, which were subsequently sliced into separate counts files based on lineage. For each lineage, batches with less than 3 cells in were removed to allow appropriate batch correction. Individual lineage files were then further subclustered and annotated with increased resolution as described above.

### TCR analysis

TCR analysis was performed using the SCIRPY package (version 0.10.1)^85^ on the T cell lineage Anndata object using default parameters.

### Gene ontology and pathway analysis

Lists of cell type-specific differentially expressed genes were generated using SCANPY’s sc.tl.rank_genes_groups function and significantly upregulated gene lists were used to generate analysis reports on the metascape web browser using the ‘Express Analysis’ function. Results were downloaded and depicted graphically using Prism (version 10).

### Deconvolution from bulk data

Published bulk RNA sequencing counts files^9^ were manually sliced according to disease and inflammatory state and deconvolved using our own single-cell RNA sequencing data using the cellanneal^86^ (v1.1.0) command line tool as per the relevant documentation. Relative percentages of cell types in each bulk dataset were then displayed graphically using Prism.

### HLA inference from single-cell RNA sequencing data

The command line tool ArcasHLA^19^ was used to infer HLA haplotypes from individual patient bam files using the extract and genotype functions. For patients PSC-2 and PSC-3 who were captured together, the filterbarcodes function in Sinto (<u>timoast.github.io/sinto</u> v0.10.1) was used to split bam files by patient before running arcasHLA extract.

### BCR Analysis

For UC and PSC-UC samples, Cell Ranger V(D)J contigs for B cells were post-processed with stand-alone IgBLAST^87^ (v1.21) aligning against the IMGT human reference directory (release 202330-1). For cells with multiple heavy or light chains the productive chain with the highest UMI count was retained. Non-productive reads, reads lacking a full length IGHV and those without a defined CDR3 were discarded from analysis. Only cells with paired heavy and light chains were used for analysis. B cell clonal lineages were built with SCOPer^88^ (v1.3.0) using single linkage for the CDR3 nucleotide sequences at a clustering threshold of 0.2 with clones defined by both heavy and light chains. For previously published healthy controls^11^, FASTQs were obtained from ArrayExpress under accession E-MTAB-9532 and processed with Cell Ranger’s *cellranger vdj* (v8.0.0). V(D)J contigs were post-processed and filtered as for UC and PSC-UC samples. Datasets were merged and summarised in R (v4.3.0) using RStudio (version 2024.04.2+764) utilising the tidyverse package^89^ (v2.0.0).

### Spatial transcriptomics data generation

STOmics *Stereo-seq Transcriptomics Set for Chip-on-a-slide user manual* (version B) was applied to optimal cutting temperature medium (OCT)-embedded fresh-frozen colonic mucosa biopsies. All samples were sectioned using the Leica CM1860 UV cryostat and two to three serial cuts were included on each Stereo-seq slide. Tissue samples were selected based on disease and inflammation status of the colon region and morphology (based on H&E). Tissue sections were adhered to the Stereo-seq chip and incubated in -20C methanol for 30 minutes. Chip IDs used in this study were D01654C4 (UC noninflamed) and D01654C5 (UC inflamed). Stereo-seq slides were fluorescently stained with Qubit ssDNA Assay Kit (Q10212; ThermoFisher) as per the manual and imaged using a Leica DM6 microscope with a CTR6 LED electronics box. The Stereo-seq *Permeabilisation Set for Chip-on-a-slide user manual* (version B) was followed to obtain an optimal permeabilisation time for colonic mucosa tissue of 8 minutes. The cDNA was purified using SPRIselect Beads (B23318; Beckman Coulter). cDNA quality was quantified before and following cDNA library preparation using Qubit 4 fluorometer with Qubit dsDNA mix (Q33230; Invitrogen) and Agilent Tapestation High Sensitivity D5000 ScreenTape Assay (5067-5592 & 5067-5593; Agilent Technologies). Four cDNA libraries were pooled to a final concentration of 13.0 ng/µL (24.63 µL volume) and sequenced on one MGI G400 FCL PE100 flow cell at the South Australian Genomics Centre.

### Spatial transcriptomics data analysis

Raw FASTQ files were processed using the Stereo-seq Analysis Workflow (SAW) pipeline (BGI). Downstream analysis was performed in Python (version 3.8.8) using Stereopy package (version 1.3.0). The data was binned to a size of 50 to combine nanopores into manageable units while preserving spatial resolution. Data was filtered by data points with gene counts 20 to 300 and mitochondrial reads less than 50%. Data was then normalised by total counts, scaled (max_value=10, zero_center=True) and log transformed. The top 1000 highly variable genes were determined using the data.tl.highly_variable_genes() function. The TACCO package (version 0.4.0.post1) was used to deconvolute spatial sequencing data using our own scRNA-seq data. The multi-centre approach with ten centres was used to account for within-cell-type variation. The ‘co_occurrence_matrix’ function was used to calculate a spatial co-occurrence score between cell types as a conditional probability for a maximum distance of 50 µm. The ggplot2 package (version 3.5.1) in R (version 4.3.3) was used to plot cell type proportion with a Wilcoxon Rank Sum Test to compare cell proportions between samples (*** for p < 0.001, ** for p < 0.01, * for p < 0.05. The likelihood of co-occurrence between cell types was visualised using the matplotlib library (version 3.8.4) in Python.

### Immunofluorescence

Immunofluorescence analysis was performed on human formalin-fixed paraffin-embedded (FFPE) tissue array slides (CO245, TissueArray.com) or on AIM study patient FFPE tissue mounted on slides as indicated in **Supplementary Table 1**. Mounted sections were stained as described previously^90^, with 30-minute antigen retrieval at pH 6. Primary and secondary antibody concentrations were as follows: EPCAM, 1:200 (ab7504, Abcam); PECAM1, 1:200 (ab76533, Abcam); C-KIT, 1:2000 (3306s, Cell Signalling); TMEM176B, 1:100 (ab236860, Abcam), goat anti-rabbit IgG AF594, 1:400 (ab150080, Abcam), goat anti-mouse IgG1 AF488, 1:400 (ab150113, Abcam). Nuclei were stained via addition of DAPI at 1:1000 during secondary antibody incubation. One drop of Prolong Diamond Antifade Mountant (P36961, ThermoFisher) was added to stained sections before mounting coverslip. Slides were imaged in a single plane on a Leica SP8 DMI6000 confocal imaging system using 29x/0.7 immersion and HC PL APO CD2 40x/1.30 oil objectives, and diode 405 nm, OPSL 488 nm, OPSL 552 nm and diode 638 nm lasers with PMT detectors. Images were processed and analysed using Imaris software (Bitplane). For quantification of IMCs, mast cells were defined using DAPI and c-KIT fluorescence, and mean AF594 fluorescence (TMEM176B) was calculated per mast cell.

### Flow cytometry

Intestinal biopsies were thawed, washed in HBSS, enzymatically digested and filtered as described above. Single cell suspensions were washed in incubation buffer (2% FCS in PBS) and split into separate microcentrifuge tubes for individual and pooled antibody staining. Cell pellets were resuspended in 100 µL incubation buffer containing individual or pooled antibodies (anti-EPCAM-BV711, 1:100, Abcam; anti-CD31-APC/Cy7, 1:100; 1µg/mL DAPI) for 30 minutes at 4°C. Stained cells were washed via centrifugation with 1 mL incubation buffer and transferred to tubes for flow cytometric analysis. Flow cytometric data was obtained using the FACS Canto II system (BD Bioscience) and analysed using FlowJo software (BD Bioscience).

### FcERI activation of mast cells

The Cl.MC/C57.1 cell line^47^ (generated from the BL6 mouse, provided by Professors John Hunt and Stephen Galli) were cultured in DMEM (high glucose and pyruvate, Gibco #11995073) supplemented with 10% FCS (SFBS, Bovogen) heat-inactivated at 56°C for 30 minutes, 1x MEM nonessential amino acids (Gibco 11140050), 2 mM L-glutamine (#25030081, Gibco) penicillin-streptomycin (#PO781, Sigma), 36 mg/L L-asparagine, 87.1 ng/L L-arginine, 6 mg/L folic acid and 50 µM 2-ME. Galli cells were suspended in complete medium at 1 million cells/mL and 15 mL was used for each condition (5 mL/replicate). For sensitisation, cells were incubated overnight with 0.1 µg DNP-IgE (Sigma D8406) per mL cell suspension. Individual samples (5 mL) were then pelleted and resuspended in 2 mL complete medium for a second, 20-minute sensitisation at 0.2 µg mouse DNP-IgE per mL. Cells were then pelleted and washed twice with PGE buffer. To induce degranulation, cells were suspended in 300 µL PGE buffer (25 mM PIPES, 120 mM NaCl, 5 mM KCl, 50 mM NaOH, 5.6 mM glucose, 0.2% BSA, 1 mM CaCl_2_) containing 100 ng/mL DNP-HSA (Sigma, #A6661) and incubated for 1 hour at 37°C with 5% CO_2_. Control samples were resuspended in PGE with no DNP-HSA. Following degranulation, 100 µL cell suspension was used to measure degranulation, and 200 µL pelleted for RNA isolation and analysis by qPCR. For BayK8644 and vehicle samples, cells were subjected to 30 minutes pre-incubation with 10 mM BayK8644 (or equivalent volume DMSO) prior to sensitisation and 10 µM BayK8644 was maintained in complete medium until degranulation.

### Assessment of mast cell degranulation

Cell suspensions were centrifuged at 300 rpm and supernatants harvested. To determine total β-hexosaminidase content, cell pellets were lysed in 100 µL 10% Triton X-100 and 25 µL of lysed cells were combined with 25 µL 5 mM P-nitrophenyl N-acetyl-B-D-glucosaminide (Sigma N9376) in 50 mM sodium citrate buffer (pH 4.5) and incubated at 37°C for 1 hour. For supernatant β-hexosaminidase content, 25 µL of supernatant was similarly combined with 25 µL 5 mM nitrophenyl and incubated at 37°C for 1 hour. After 1 hour, reactions were terminated with 250 µL 0.2 M glycine-NaOH (pH 10.6) and absorbance at 405 nm determined using a Nanodrop. Percentage β-hexosaminidase (βhex) release was calculated by the equation:

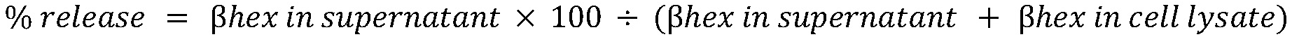

### TLR4 activation of mast cells

Staphylococcus aureus particles (S2859, Invitrogen) were suspended in PBS at 1 million/µL. Galli cells were suspended at 1 million/mL in complete medium and 3 mL cell suspension (3 million cells) seeded per well into 6-well plates. Four conditions were studied: Activation only (addition of 15 million staphylococcus beads), negative control (addition of 15 µL PBS only), activation in presence of vehicle (addition of 3 µL DMSO followed by 15 million staphylococcus beads after 30 minutes), and activation in presence of TMEM176b inhibitor BayK8644 (10 mM BayK8644 followed by 15 million staphylococcus beads after 30 minutes). For each condition, four timepoints were assessed: 0 hours (pre-activation or after incubation with vehicle/BayK8644), 4 hours post-activation, 12 hours post-activation, and 24 hours post-activation. Each condition and time point was assessed in triplicate. At each timepoint, cells were centrifuged for 5 minutes at 300 g and supernatant discarded. Cell pellets were lysed in 350 µL buffer RLT (QIAgen) and immediately stored at -80C.

### Quantitative real-time PCR

RNA was isolated from fresh or thawed cell lysates using the RNeasy Mini Kit (QIAgen) as per manufacturer’s instructions. RNA concentrations were measured via Nanodrop and cDNA generated from 2 µg RNA using the High Capacity cDNA Reverse Transcription Kit (#4368814, ThermoFisher Scientific) as per manufacturer’s instructions. RT-qPCR was performed in 10 µL reactions containing 2 µg cDNA, 5 µL 2X HotStart PCR Master Mix (HMM, MCLAB), 0.7 µL SYTO9 and mouse primer pairs topped up with nuclease-free H_2_O. Primer information and concentrations are shown in **Table 1**. Reactions were analysed in a LightCycler 480 System (Roche) with 15 minute heat inactivation at 95°C followed by 45 PCR cycles (15 seconds at 95°C, 30 seconds at 58°C, 15 seconds at 72°C).

**Table 1.**
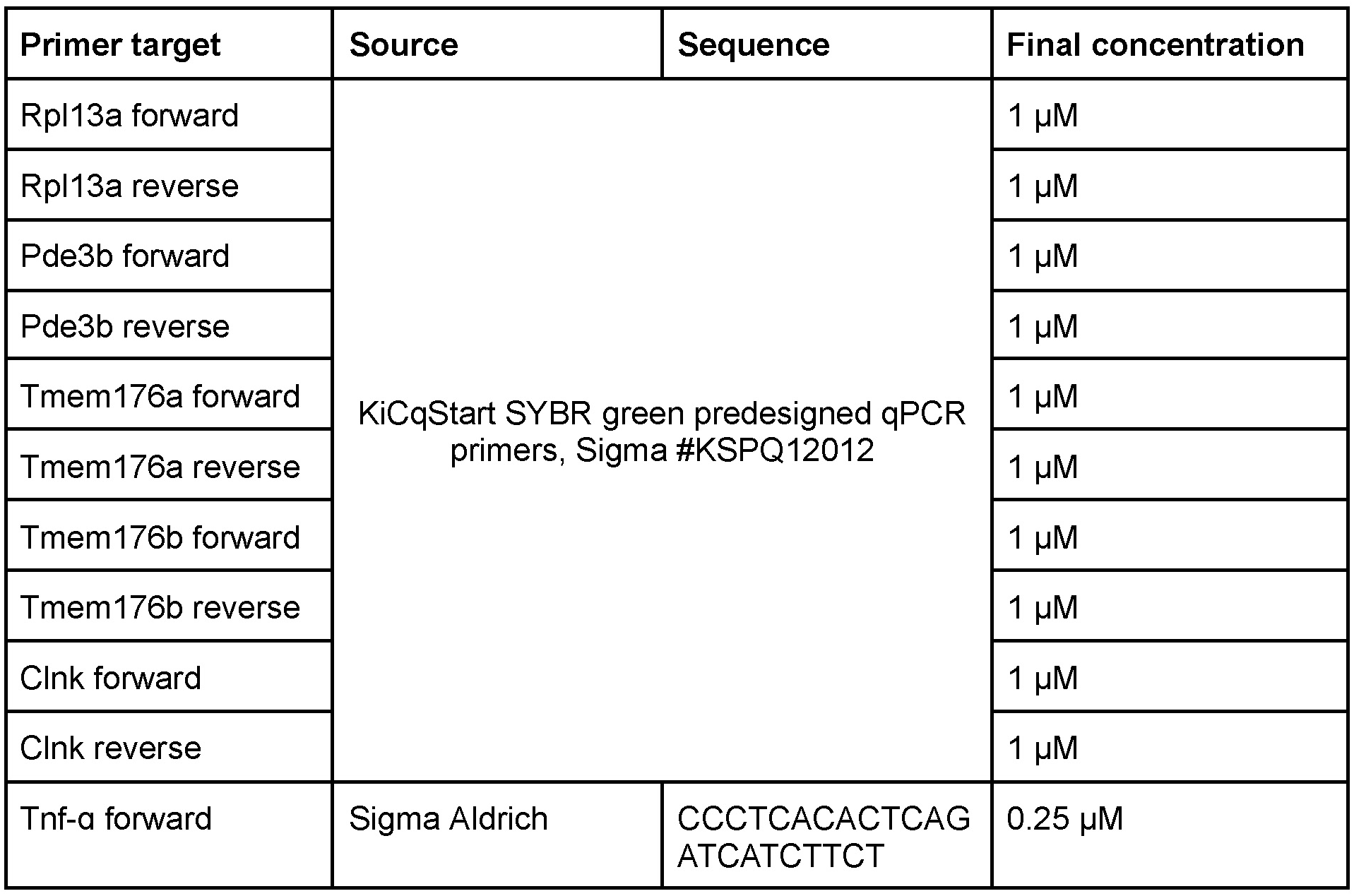
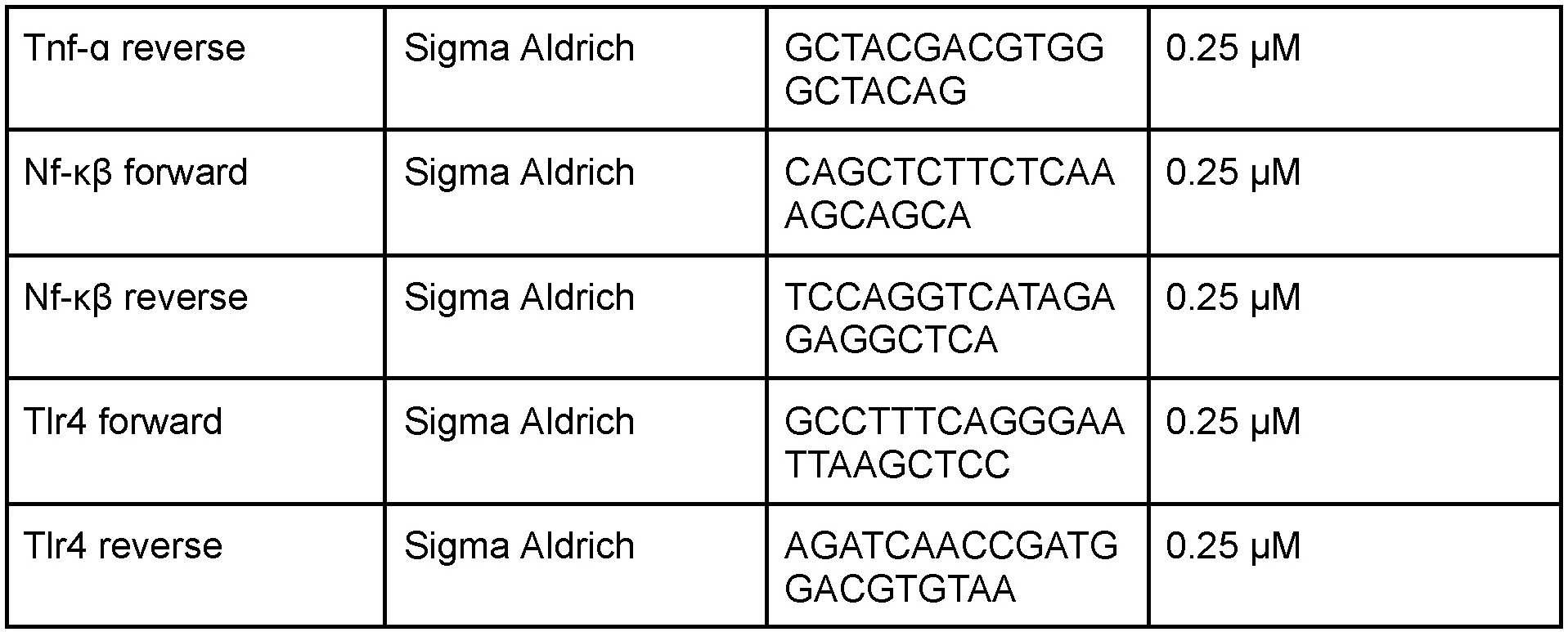
Primer pairs.

### Data availability

Raw single-cell RNA sequencing data files will be available through ArrayExpress following peer-review. Sequencing data for the microbiome will be available at MGnify following peer-review.

## Acknowledgments

We acknowledge the support received from the Garvan Cellular Genomics, Histology, Microscopy, and Molecular Genetics platforms. We sincerely thank W. Muskovic for his critical help with troubleshooting data analysis. We thank J.E. Powell and C. Chaffer for insightful discussions about the project. We acknowledge technical support provided by the Garvan Cellular Genomics and Data Science Platforms, and Tissue Culture and Microscopy Core Facilities. This research was supported by funding from the Spinak Fellowship (K.R.J), a National Health and Medical Research Investigator Fellowship (grant no. APP1194063 to K.R.J), a Ramaciotti Health Investment grant (2021HIG72 to K.R.J) and Perpetual IMPACT Philanthropy grant (PR10810 to K.R.J). We sincerely thank and acknowledge the donors who are part of the Australian IBD Microbiome (AIM) Study for access to their clinical specimens.

## Author contribution

K.R.J. and S.G. designed the study. S.G., A.K., P.T. and P.S. coordinated patient recruitment, consent and biospecimen collection. J.L.E.T., K.R.J. and S.K. carried out scRNAseq and 16S rDNA sequencing data generation. Analysis of the 16S rDNA sequencing was performed by J.L.E.T. and F.Z. Analysis of scRNA-seq data was performed by J.L.E.T. and K.R.J. Analysis of the BCR sequencing data was carried out by K.J.L.J. J.W. sectioned fresh frozen tissue for spatial transcriptomics. J.L.E.T. carried out microscopy, flow cytometry and validation experiments. Spatial transcriptomics data generation was performed by K.R.J. and P.M. Analysis of the spatial transcriptomics data was performed by P.M. with support from C.W. and A.H. G.L.H. oversaw and provided reagents for experiments relating to the microbiome. N.T. oversaw and provided reagents for experiments relating to mast cells. H.W.K. oversaw scRNAseq data analysis and provided data interpretation. J.L.E.T. and K.R.J. wrote the manuscript. All authors provided critical review of the manuscript.

---

**Supplementary Figure 1.**
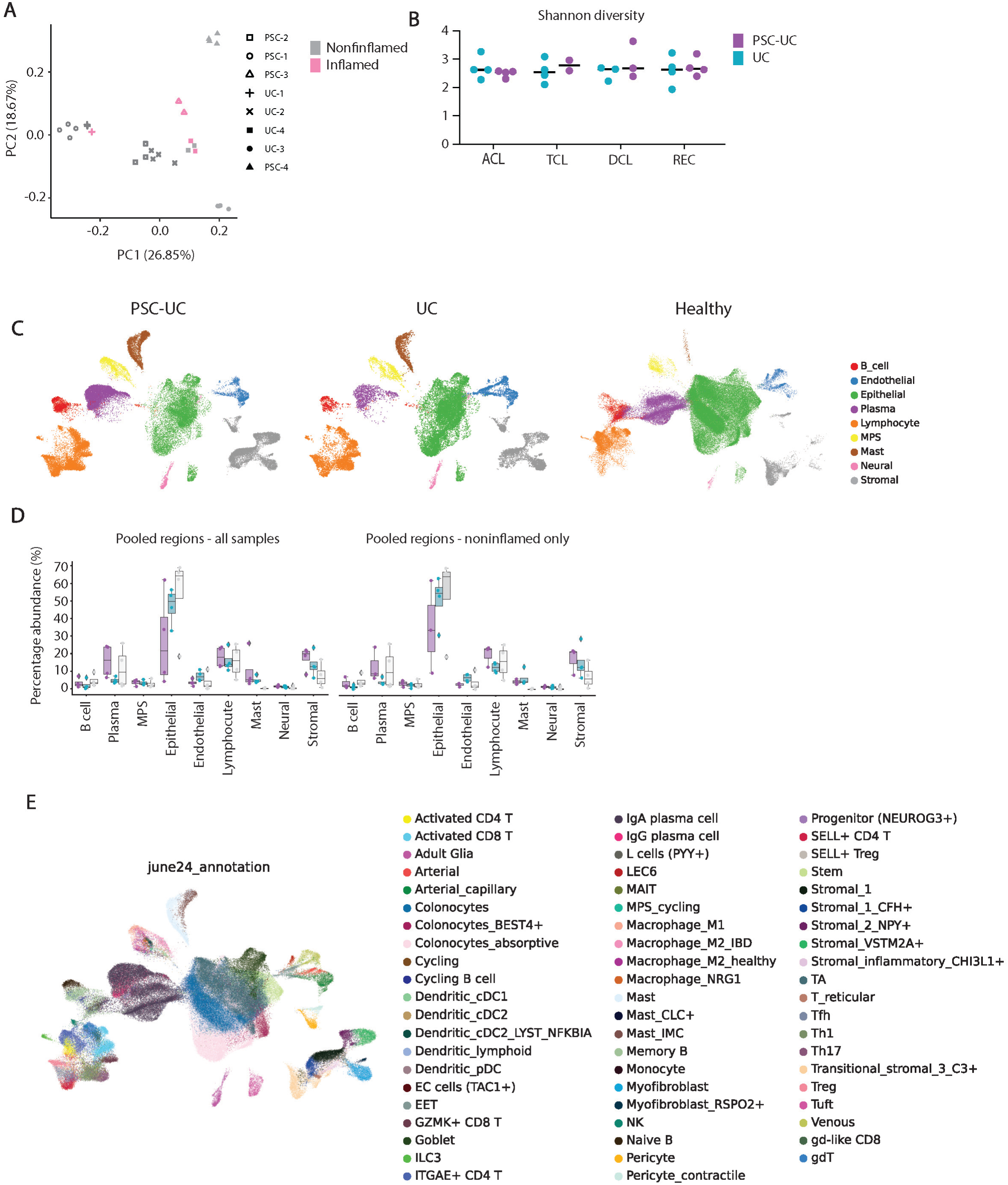
**A |** Principal coordinate analysis (PCoA) plot of 16S rRNA gene sequencing of colonic mucosa biopsies from the ascending, transverse, descending colon and rectum of patients with UC and PSC-UC (n=4/patient), with inflammation state of tissue denoted by colour and patient denoted by symbols. **B |** Shannon diversity dot plot of colonic microbiome for PSC-UC and UC patients per region. **C |** UMAPs of the entire scRNA-sequencing dataset separated by disease status and coloured by broad cell lineage. HC healthy control^11^, UC ulcerative colitis, PSC-UC primary sclerosing cholangitis concomitant with colonic colitis. **D |** Percentage abundance of cell lineages per patient for all cells (left) and excluding cells mapped to inflamed regions (right). **E |** UMAP of the full scRNA-sequencing dataset of HC, UC and PSC-UC patients coloured by cell type annotation.

**Supplementary Figure 2.**
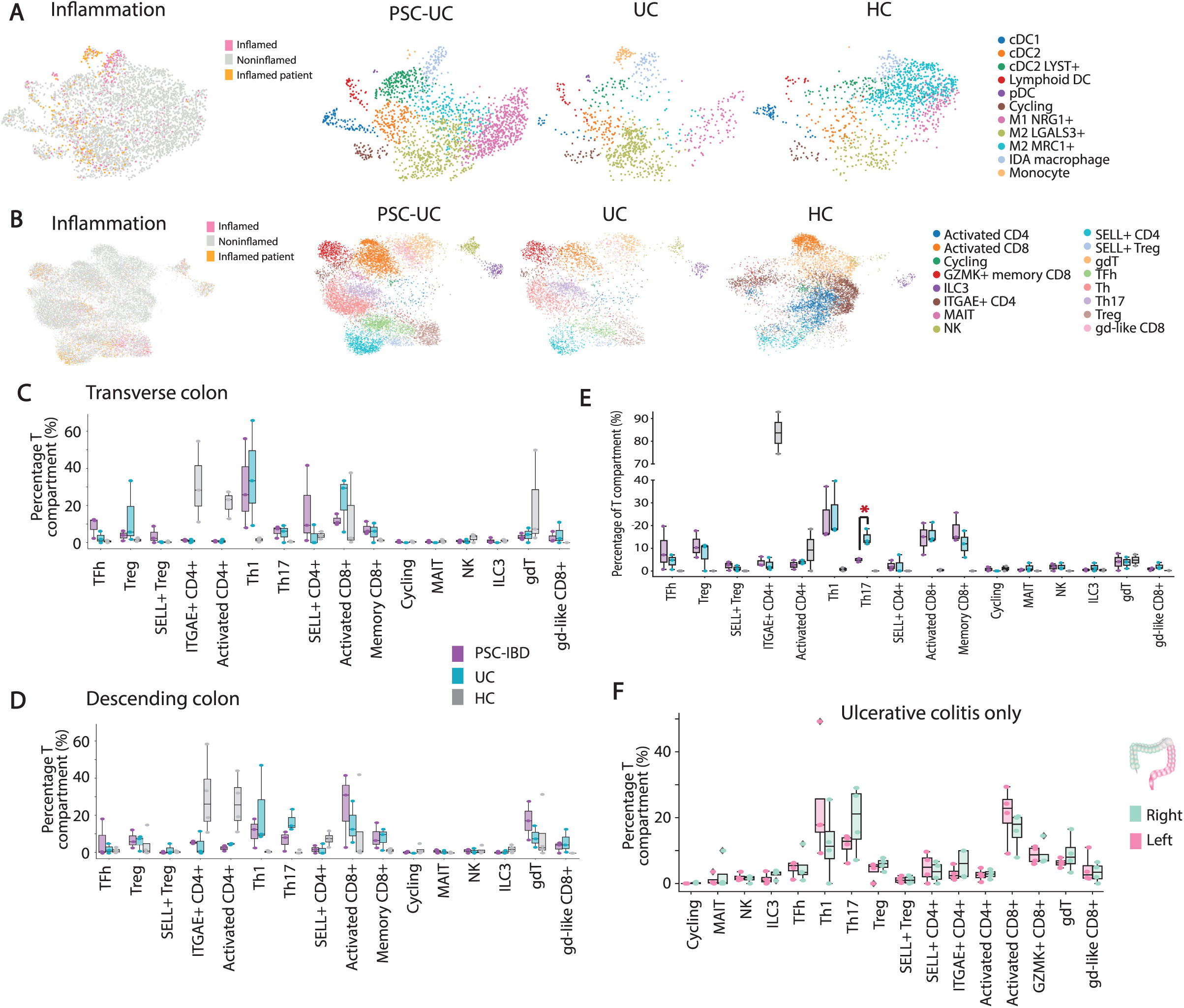
**A |** UMAP of the scRNA-sequencing dataset mononuclear phagocyte system (MPS) compartment coloured by inflammation (left) and separated by disease status (right) and coloured by cell type annotation. UC ulcerative colitis, PSC-UC primary sclerosing cholangitis concomitant with colonic colitis. HC healthy control. **B |** UMAP of the T cell compartment coloured by inflammation (left) and separated by disease status coloured by cell type annotation. Th, T helper; Treg, T regulatory; ILC, innate lymphoid cell; NK, natural killer cell; MAIT, mucosal-associated invariant T cell. **C |** Relative abundance analyses for the T cell compartment in the transverse colon. Each dot represents one biopsy. **D |** Relative abundance analyses for the T cell compartment in the descending colon. Each dot represents one biopsy. **E |** Relative abundance analyses for the T cell compartment in the rectum. Each dot represents one biopsy. P value for PSC-UC vs. UC comparisons as determined by unpaired T test = 0.0101 for Th17. **F |** Relative abundance of T cell subtypes in the right (pooled ascending and transverse biopsies) and left (pooled descending and rectum biopsies) colons of ulcerative colitis patients. For all abundance analyses, cells mapping to inflamed regions were excluded prior to analysis.

**Supplementary Figure 3.**
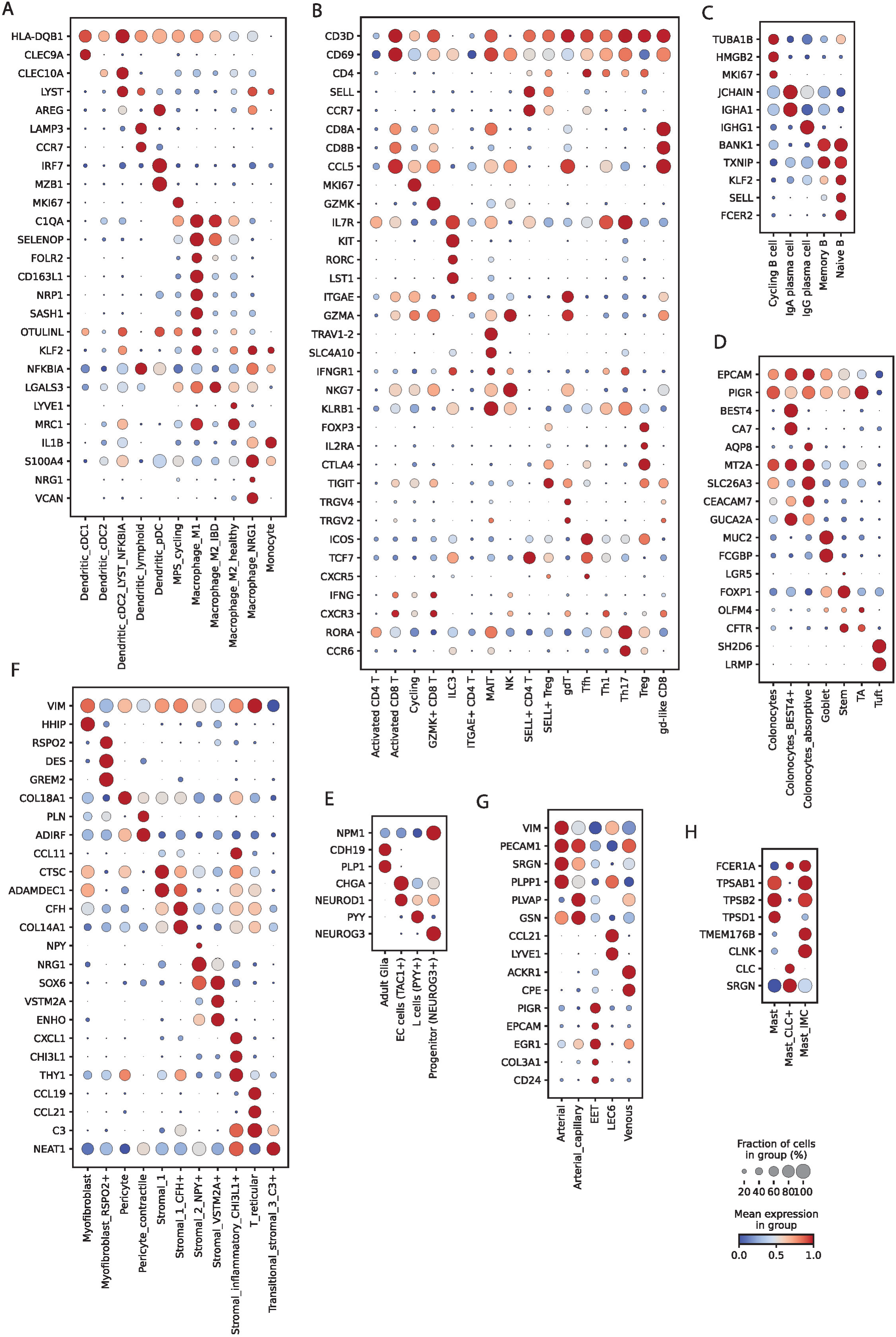
Gene expression of marker genes for each compartment of the scRNA-sequencing dataset. Mononuclear phagocyte system (**A**), T compartment (**B**), B and plasma cells (**C**), epithelial compartment (**D**), neural compartment (**E**), stromal compartment (**F**), endothelial compartment (**G**) and mast cells (**H**).

**Supplementary Figure 4.**
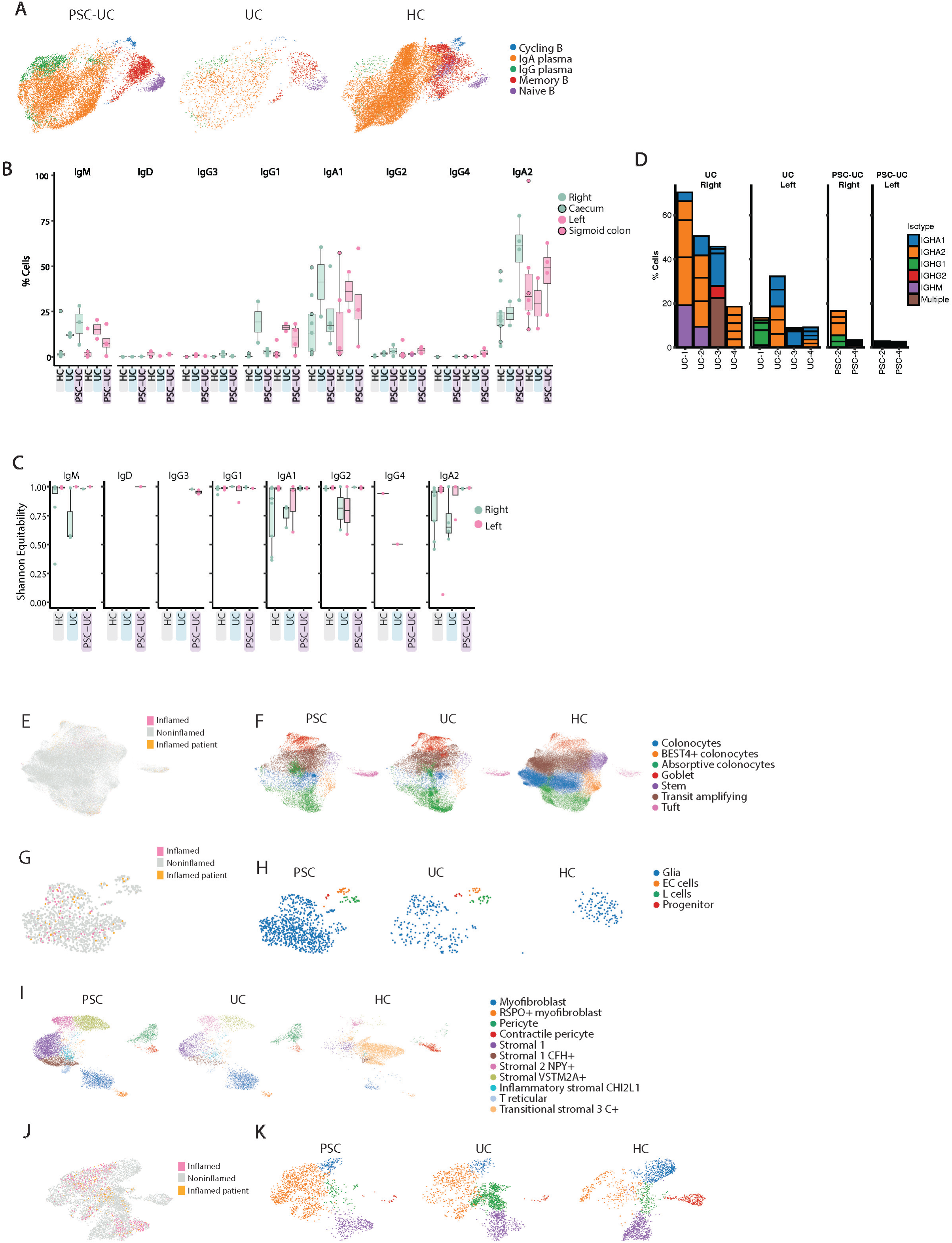
**A** | UMAP of the subsetted B lineage cells from scRNA-sequencing data separated by disease state and coloured by cell type annotation. **B** | Percent abundance of each isotype class within total functional B cell receptor (BCR) sequences from scRNA-sequencing data of healthy controls (HC) and UC and PSC-UC patients. Colour points represent pooled colonic biopsies from right (ascending or transverse colon) or left (descending colon or rectum) colon. Outline further denotes caecum (green) and sigmoid colon (pink) of HCs. Only sequences from noninflamed colon regions are included. **C** | Shannon Equitability scores of BCR sequences separated by isotype of noninflamed colon regions of UC and PSC-UC patients. Colour points represent pooled colonic biopsies from right (ascending and transverse colon) or left (descending colon and rectum) colon. ANOVA (cohort + side + isotype) with Tukey HSD post-test and multi hypothesis corrections showed no significance. **D** | Percentage of total BCR sequences that are accounted for by top five expanded clones of noninflamed colon regions from UC and PSC-UC patients (x-axis) coloured by heavy chain usage (isotype). ‘Multiple’ refers to clonal groups with members of multiple isotypes. **E** | UMAP of scRNA-sequencing of epithelial lineage coloured by colonic mucosa inflammatory state and **F** | annotated celltype separated by cohort. **G** | UMAP of scRNA-sequencing dataset neural lineage compartment coloured by colonic mucosa inflammatory state and **H** | annotated celltype separated by cohort. **I** | UMAP of scRNA-sequencing dataset stromal lineage compartment coloured by annotated celltype separated by cohort. **J** | UMAP of scRNA-sequencing of endothelial lineage compartment coloured by colonic mucosa inflammatory state and **K** | Leiden clustering separated by cohort. ‘Inflamed patient’ in **E**, **G** and **J** indicates a patient experiencing inflammation at the time of tissue sampling but hashtag oligonucleotides were unable to confidently separate cells by colon regions so exact inflammatory state can not be determined.

**Supplementary Figure 5.**
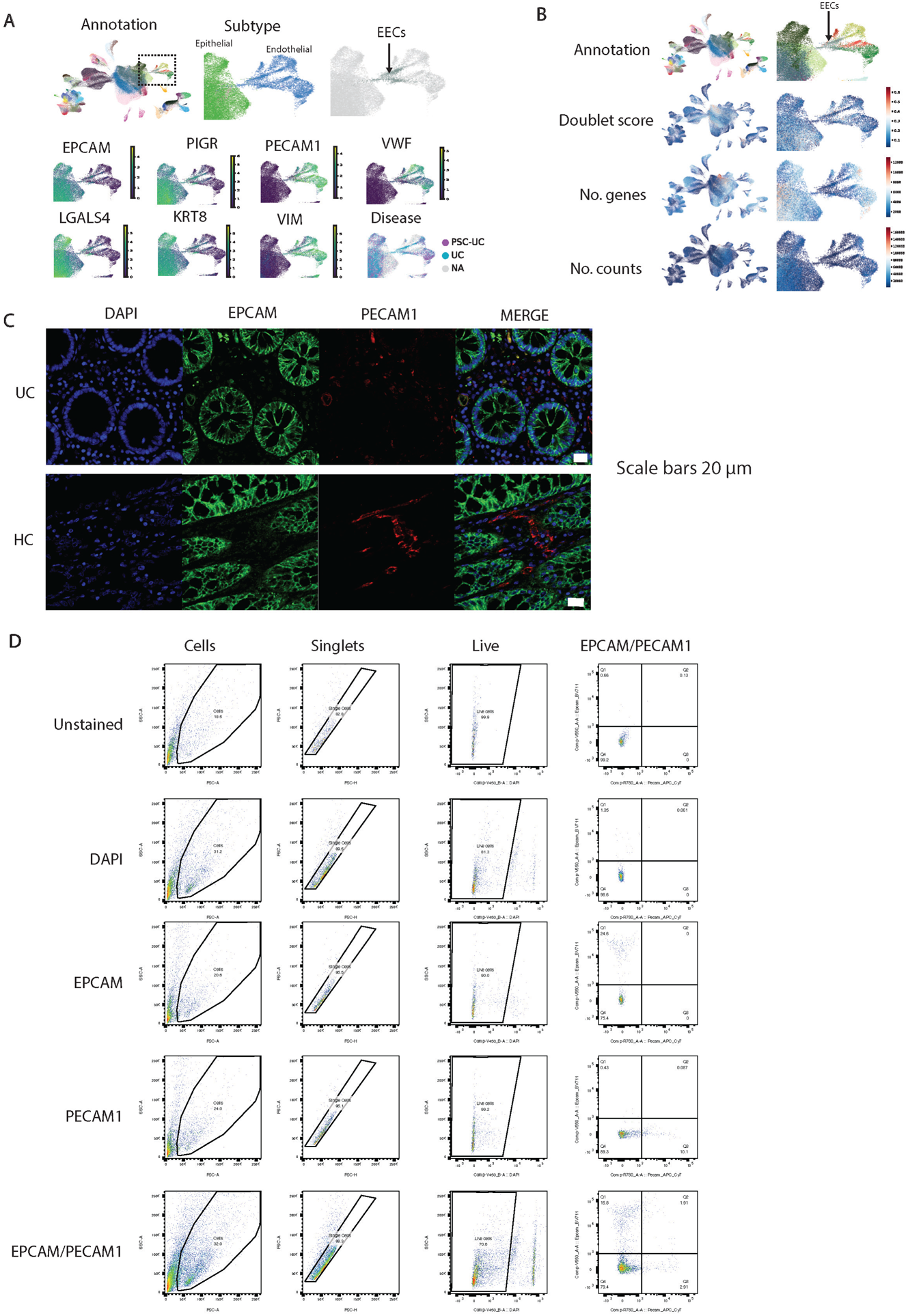
**A |** UMAP of the entire scRNA-sequencing dataset (left) coloured by annotation and indicating region of interest containing epithelial (green) and endothelial broad cell lineages (blue). Right shows close-up of the region of interest coloured by cell type annotation with EECs in grey. Further UMAPs show individual gene expression and disease state. **B |** UMAPs of the entire scRNA-sequencing dataset (left) coloured by cell type annotation, doublet score, number of genes and number of counts per cell. **C |** Immunofluorescence images showing individual DAPI, EPCAM and PECAM1 fluorescence in ulcerative colitis (UC) and healthy controls (HC). Scale bars show 20 µm. **D |** Flow cytometry gating strategies in unstained, single stained and fully-stained UC patient sample.

**Supplementary Figure 6.**
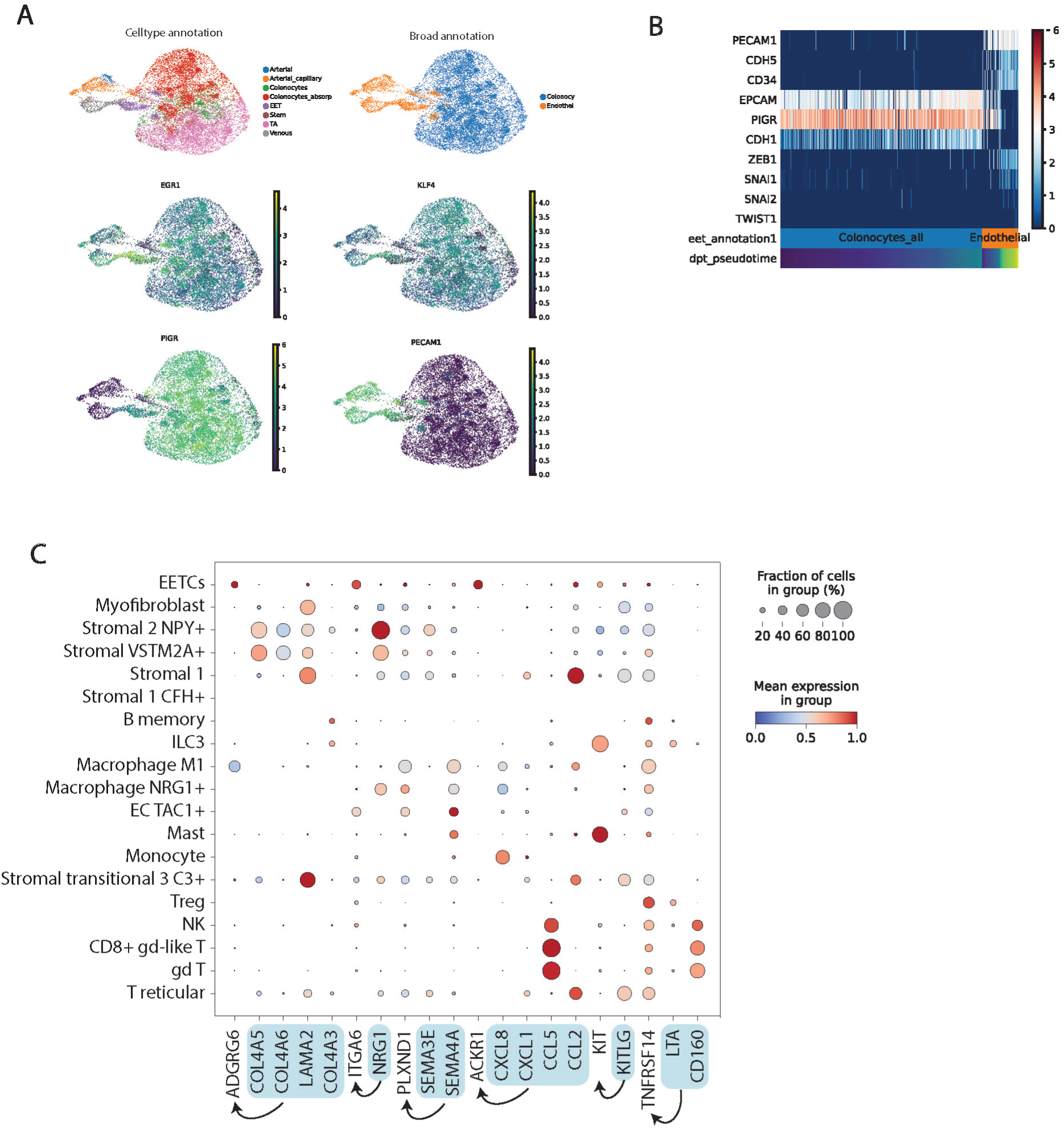
**A |** UMAP of the scRNA-sequencing dataset subsetted to include arterial, arterial capillary, colonocytes, absorptive colonocytes, epithelial-endothelial cells (EECs), stem cells, transit amplifying (TA) cells and venous endothelial cells coloured by subtype (top left), broad cell type annotation (top right) and by EGR1, KLF4, PIGR and PECAM1 gene expression. **B |** Expression of endothelial (PECAM1, CDH5, CD34), epithelial (EPCAM, PIGR, CDH1) and major epithelial-mesenchymal transition genes of cells in A ordered by pseudotime. **C |** Dot plot showing expression of enriched transcripts in EECs and interacting partners identified by CellphoneDB.

**Supplementary Figure 7.**
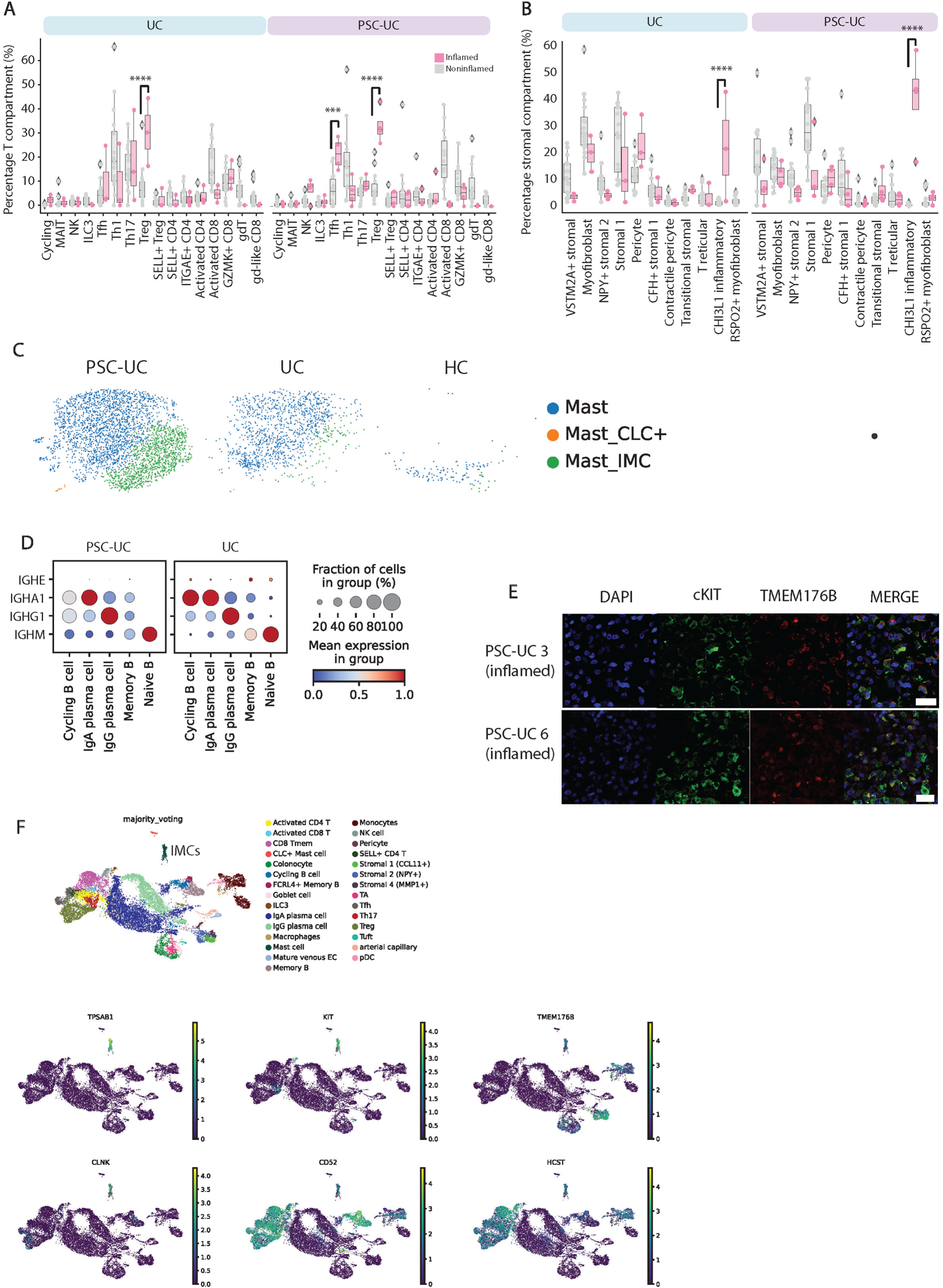
**A |** Percentage of T cell subtypes within the T compartment for inflamed versus noninflamed samples. Each dot represents one biopsy. P value for Tfh PSC = 0.001, Treg in UC = 0.0009, Treg in PSC-UC = 2.6×10^-6^ as determined by unpaired T test. **B |** Percentage of stromal subtypes within the stromal compartment for inflamed versus noninflamed samples. Each dot represents one biopsy. P value for CHI3L1+ inflammatory fibroblasts in UC = 0.0002 and in PSC = 5.9×10^-11^ as determined by unpaired T test. **C |** UMAP of the scRNA-sequencing dataset mast cell compartment coloured by cell type annotation and separated by disease status. HC, healthy control^11^; UC, ulcerative colitis; PSC-UC, primary sclerosing cholangitis concomitant with colitis. **D |** Dot plot showing expression of immunoglobulin transcripts in B and plasma cells in noninflamed PSC-UC and UC patient biopsies. **E |** Immunofluorescence imaging of KIT+ TMEM176B+ mast cells in PSC-UC

